# Single cell transcriptional evolution of myeloid leukaemia of Down syndrome

**DOI:** 10.1101/2025.04.01.646181

**Authors:** Mi K. Trinh, Konstantin Schuschel, Hasan Issa, Rebecca Thomas, Conor Parks, Agnes Oszlanczi, Toochi Ogbonnah, Di Zhou, Lira Mamanova, Elena Prigmore, Emilia Robertson, Angus Hodder, Anna Wenger, Nathaniel D. Anderson, Holly J. Whitfield, Taryn D. Treger, José Gonçalves-Dias, Karin Straathof, David O’Connor, Matthew D. Young, Laura Jardine, Stuart Adams, Jan-Henning Klusmann, Jack Bartram, Sam Behjati

**Affiliations:** Wellcome Sanger Institute; Hinxton, CB10 1SA, UK; Department of Pediatrics, Goethe University Frankfurt, Frankfurt, Germany; Frankfurt Cancer Institute, Frankfurt am Main, Germany; Great Ormond Street Hospital for Children NHS Foundation Trust, London, WC1N 3JH, UK; Department of Paediatrics, University of Cambridge, Cambridge, CB2 0QQ, UK; Cambridge University Hospitals NHS Foundation Trust, Cambridge, CB2 0QQ, UK; UCL Cancer Institute, 72 Huntley St, London, WC1E 6DD, UK; Great Ormond Street Biomedical Research Centre, 30 Guilford Street, London, WC1N 1EH, UK; UCL Great Ormond Street Institute of Child Health; London, WC1N 1EH, UK; Biosciences Institute, Newcastle University; Newcastle upon Tyne, NE2 4HH, UK; German Cancer Consortium (DKTK), Partner Site Frankfurt/Mainz and German Cancer Research Center (DKFZ), Heidelberg, Germany

## Abstract

Children with Down syndrome have a 150-fold increased risk of developing myeloid leukaemia (ML-DS). Unusually for a childhood leukaemia, ML-DS arises from a preleukaemic state, termed transient abnormal myelopoiesis (TAM), via a conserved sequence of mutations. Here, we examined the relationship between the genetic and transcriptional evolution of ML-DS from natural variation; a rich collection of primary patient samples and fetal tissues with a range of constitutional karyotypes. We distilled transcriptional consequences of each genetic step in ML-DS evolution, utilising single cell mRNA sequencing, complemented by phylogenetic analyses in progressive disease. We found that transcriptional changes induced by the TAM-defining *GATA1* mutations are retained in, and account for most of the ML-DS transcriptome. The *GATA1* transcriptome pervaded all stages of ML-DS, including progressive disease that had undergone genetic evolution. Our approach delineates the transcriptional evolution of ML-DS and provides an analytical blueprint for distilling consequences of mutations within their pathophysiological context.

## INTRODUCTION

Children with constitutional trisomy of chromosome 21 (i.e. Down syndrome) have a 150-fold increased risk of developing myeloid leukaemia^1–3^. Myeloid leukaemia of Down syndrome (ML-DS) evolves from a preleukaemic state, termed transient abnormal myelopoiesis (TAM), although it may appear and resolve unnoticed in the postnatal period. Approximately 15%-30% of children with Down syndrome have molecular evidence of TAM at birth^4,5^. Morphologically, the leukaemic cells (blasts) of ML-DS and TAM are indistinguishable^6^, exhibiting megakaryoblastic differentiation and immunophenotypic features (e.g. CD41, CD61 and lineage-atypical features e.g. CD7, CD56.^7^). There are no distinctive diagnostic features that separate TAM from ML-DS. The key difference lies in the clinical trajectory of these diseases. ML-DS is invariably fatal if left untreated; it is cancer. By contrast, TAM is a self-limiting clonal expansion. It only requires treatment when resolution needs to be hastened because of significant physiological perturbations^8,9^. In clinical practice, age cut-offs are applied to distinguish between TAM and ML-DS, with the former being confined to the first few months of life (6 months in UK practice (http://www.cclg.org.uk/, Tunstall *et al.*, 2018^9^)).

ML-DS has been extensively studied. As a stepwise model of leukaemogenesis, it may provide insights generalisable to other leukaemia subtypes where a precursor phase cannot be examined. In particular, the genetic basis of TAM and ML-DS has been resolved in detail **Figure 1A**). The first step towards transformation is constitutional trisomy 21. It has been shown to bias fetal haematopoiesis towards erythroid and megakaryocytic lineages at the cost of the B cell lineage^10–13^. This is followed by the acquisition of oncogenic mutations in the *GATA1* gene which underpin the development of TAM^14–16^. GATA1 is a transcription factor that plays a critical role in regulating erythroid and megakaryocytic fate, via upregulation of genes such as *FOG1* and *KLF1* and downregulation of genes including *KIT, MYC* and *GATA2*^17,18^. Mutations in TAM yield a truncated GATA1 protein, which permits some differentiation, but fails to adequately suppress pro-proliferative features^19,20^. In addition to *GATA1* mutations, ML-DS harbours cancer causing (“driver”) variants in 75%-93% of cases^1,21^, commonly affecting genes encoding JAK kinases, the cohesin complex, or epigenetic regulators. Between 7%-25% of ML-DS cases do not acquire further discernible driver events, yet clinically represent a malignant disease rather than a self-resolving clonal expansion of blood. Extensive phenotypic, genetic and detailed modelling work notwithstanding, the effects of each leukaemogenic event in the evolution of ML-DS remains incompletely understood^16,22^.

**Figure 1:**
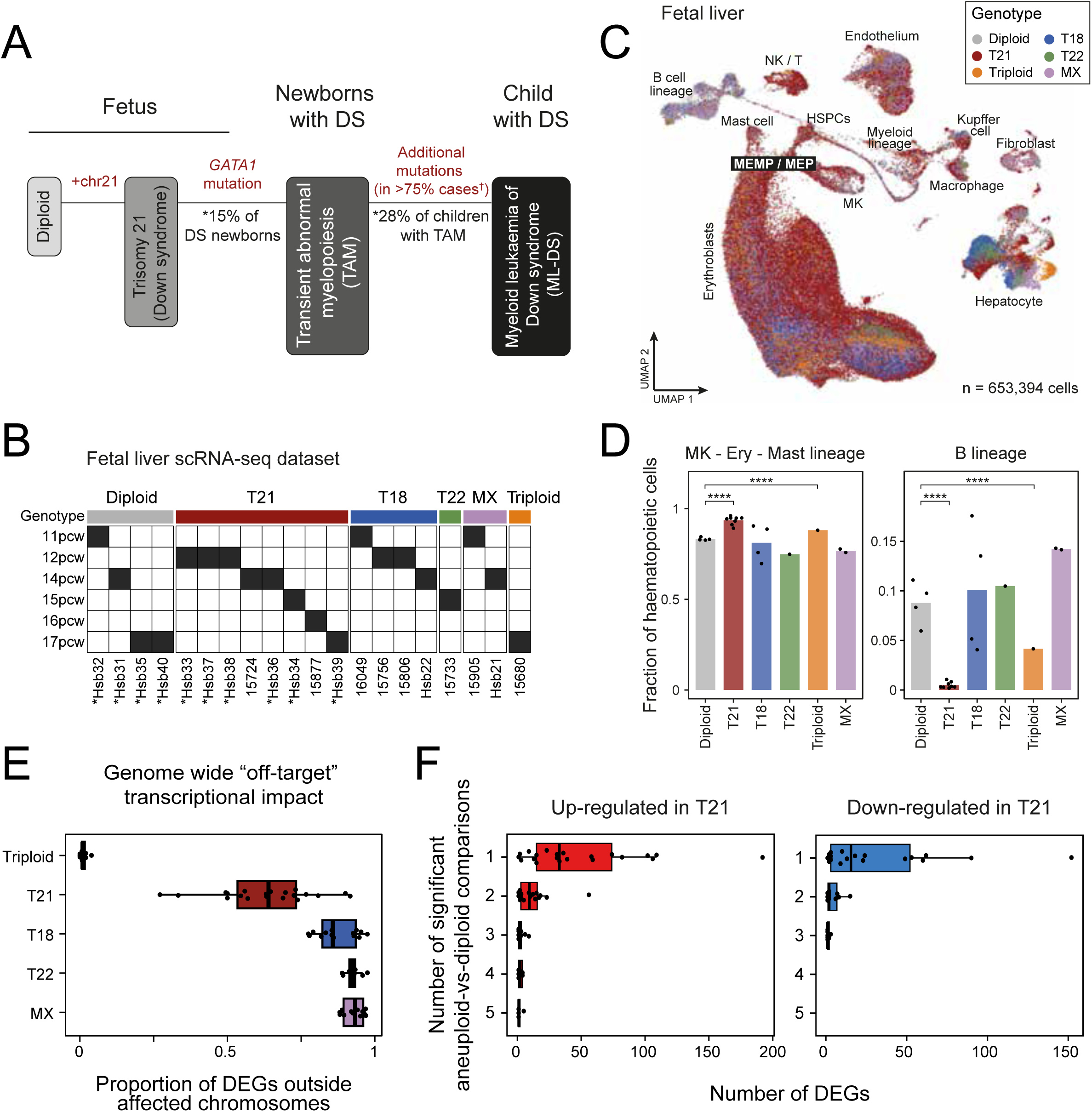
Transcriptional effects of karyotypic variants on fetal hepatic haematopoiesis. (A) Schematic of the stepwise pathogenesis of ML-DS. Statistics were obtained from: (*) Goemans *et al*., 2021^5^ for the frequency of *GATA1* mutations in neonates with Down syndrome (DS); and (†) Labuhn *et al.*, 2019^1^ and Sato *et al.*, 2024^21^ for the number of ML-DS cases with known additional driver mutations. (B) Fetal liver samples from fetuses with different karyotypes. Asterisks (*) indicate samples where cells were sorted into CD45+ and CD45-fractions before sequencing. (C) Uniform Manifold Approximation and Projection (UMAP) visualisation of the fetal liver scRNA-seq dataset, with cells (dots) coloured by genotype. (D) Bar plot showing the mean proportion of megakaryocyte-erythroid-mast cell compartment and B lineage among all haematopoietic cells across donors for each genotype. Dots represent individual fetal samples within the genotype group. Asterisks (*) indicate significantly higher megakaryocyte-erythroid-mast cell proportions and lower B lineage proportions in trisomy 21 and triploid samples compared to diploid samples (p-value < 2.2*10^-16^, one-tailed two-proportion z-test). (E) Box plot showing the impact of different karyotypes on genome-wide gene expression across cell types (dots) compared to diploid samples, with the genome-wide “off-target” transcriptional impact defined as the proportion of differentially expressed genes (DEGs) residing outside the affected chromosome(s). (F) Number of DEGs identified in each trisomy 21 cell type (dots) when compared to their diploid counterparts. DEGs are further categorised based on whether they were also detected as differentially expressed in one or more other abnormal karyotypes (indicated by the number of comparisons - x axis). ***Abbreviation*** DS - Down syndrome. pcw - post conception week. <u>Cell types</u>: HSPCs - haematopoietic stem and progenitor cells; CMP / GMP - common myeloid progenitor / granulocyte-monocyte progenitor; DC - dendritic cell; Ery - erythroblast; MEMP / MEP - megakaryocyte-erythroid-mast cell progenitor / megakaryocyte-erythroid progenitor; MK - megakaryocyte; NK / T - natural killer cell / T cell; pDC - plasmacytoid dendritic cell. <u>Genotypes</u>: T - trisomy; MX - monosomy X.

The advent of single cell transcriptomics provides an opportunity to directly distil cellular consequences of mutations from natural variation. It delivers with every cell an independent readout of an individual’s disease, whilst studying the same condition across patients enables the identification of overarching signals. The key advantage of this approach is that it examines variants within their physiological context, rather than in the engineered environment of model systems. A central limitation is the availability of adequate patient samples (fresh material or viably frozen cells), especially in rare disease such as TAM and ML-DS.

We set out to distil the cellular difference between TAM and ML-DS from natural variation, by reconstructing the transcriptional evolution of ML-DS via trisomy 21 and TAM. Harnessing tissue banking efforts of the UK’s largest children’s cancer unit and a German cohort from an international ML-DS study, we were able to examine the entire spectrum of TAM and ML-DS directly from patient samples, including rare cases of refractory and recurrent disease, in the context of human fetal tissues and other relevant leukaemias.

## RESULTS

### Study cohort overview

We assembled a cohort of primary patient samples from the UK and Germany encompassing TAM and ML-DS, including rare cases of recurrent TAM, recurrent ML-DS, and refractory ML-DS (**Table 1, Supplementary Table 1**). All children were younger than 5 years of age at ML-DS diagnosis, and all cases had megakaryoblastic differentiation on flow cytometry profiling. In addition, we studied other leukaemias to disentangle the effects of mutations, cell lineage, and phenotype on transcription: an exceptional rare case of diploid myelodysplastic syndrome (MDS) harbouring a *GATA1* mutation, as well as a case of MDS without *GATA1* mutation; acute megakaryocytic leukaemia (AMKL) in disomic children (n=2); other childhood myeloid leukaemias (AML, n=5) to capture myeloid cancer phenotypes; B cell lymphoblastic leukaemias (B-ALL) arising in children with (n=8) or without Down syndrome (n=2), to assess the effects of trisomy 21 in a different haematopoietic line; and a solid haematological B cell neoplasm arising in a child with Down syndrome (central nervous system lymphoproliferative disorder; LPD). To study the effects of trisomy 21 on fetal haematopoiesis, we examined fetal livers (thought to be the origin of preleukaemic cells in Down syndrome^23^) from diploid donors and donors with variant constitutional karyotypes (trisomy 21, trisomy 18, trisomy 22, monosomy X and whole genome trisomy or triploidy; **Supplementary Table 1**). All samples (fresh or viably frozen) were enriched for live cells, prior to single cell mRNA sequencing (scRNA-seq) using the Chromium 10X platform. We processed the scRNA-seq data in a standardised way, including removal of ambient mRNAs and doublets, to derive high quality count tables of transcripts for 653,394 fetal liver cells, and 525,650 patient cells: 196,090 normal and 329,560 neoplastic cells. In addition, we performed whole genome sequencing where possible to enable integrated analyses of transcriptional and genetic evolution, as detailed in individual sections. A detailed account of the study cohort including assays and data volume is provided in **Supplementary Table 1**.

**Table 1.**
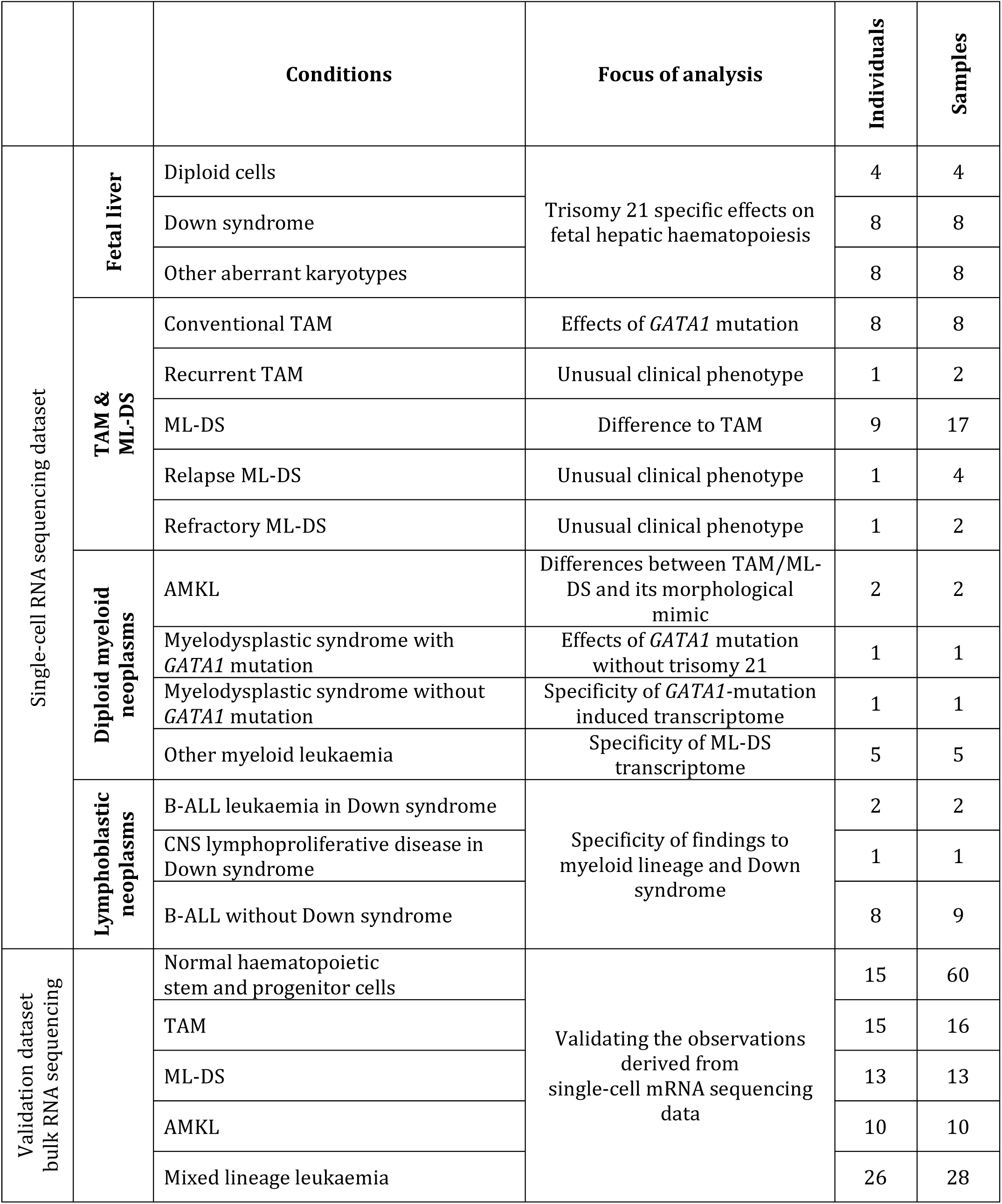
Overview of study cohort.

|  |  | Conditions | Focus of analysis | Individuals | Samples |
| --- | --- | --- | --- | --- | --- |
| Single-cell RNA sequencing dataset | Fetal liver | Diploid cells | Trisomy 21 specific effects on fetal hepatic haematopoiesis | 4 | 4 |
|  |  | Down syndrome |  | 8 | 8 |
|  |  | Other aberrant karyotypes |  | 8 | 8 |
|  | TAM & ML-DS | Conventional TAM | Effects of *GATA1* mutation | 8 | 8 |
|  |  | Recurrent TAM | Unusual clinical phenotype | 1 | 2 |
|  |  | ML-DS | Difference to TAM | 9 | 17 |
|  |  | Relapse ML-DS | Unusual clinical phenotype | 1 | 4 |
|  |  | Refractory ML-DS | Unusual clinical phenotype | 1 | 2 |
|  | Diploid myeloid neoplasms | AMKL | Differences between TAM/ML-DS and its morphological mimic | 2 | 2 |
|  |  | Myelodysplastic syndrome with *GATA1* mutation | Effects of *GATA1* mutation without trisomy 21 | 1 | 1 |
|  |  | Myelodysplastic syndrome without *GATA1* mutation | Specificity of *GATA1*-mutation induced transcriptome | 1 | 1 |
|  |  | Other myeloid leukaemia | Specificity of ML-DS transcriptome | 5 | 5 |
|  | Lymphoblastic neoplasms | B-ALL leukaemia in Down syndrome | Specificity of findings to myeloid lineage and Down syndrome | 2 | 2 |
|  |  | CNS lymphoproliferative disease in Down syndrome |  | 1 | 1 |
|  |  | B-ALL without Down syndrome |  | 8 | 9 |
| Validation dataset bulk RNA sequencing |  | Normal haematopoietic stem and progenitor cells | Validating the observations derived from single-cell mRNA sequencing data | 15 | 60 |
|  |  | TAM |  | 15 | 16 |
|  |  | ML-DS |  | 13 | 13 |
|  |  | AMKL |  | 10 | 10 |
|  |  | Mixed lineage leukaemia |  | 26 | 28 |

### Effects of karyotypic variants on fetal hepatic haematopoiesis

To establish the contribution of trisomy 21 to the ML-DS transcriptome, we compared liver haematopoietic cells of fetuses with trisomy 21 against disomic fetal liver cells. We expanded our analyses to liver haematopoietic cells from fetuses with a range of karyotypic variants – trisomy 18, trisomy 22, monosomy X and triploidy (**Figure 1B, Supplementary Table 1**) – to establish whether the changes induced by trisomy 21 are specific to Down syndrome or represent generic perturbation of haematopoiesis caused by numerical variation of the human karyotype. Overall, our fetal liver scRNA-seq dataset captured 31 different cell types across both haematopoietic and stromal compartments, with representation of all genotypes (**Figure 1C**, **Supplementary Figure 1**). In keeping with previous work^10,11,13^, we found that in trisomy 21, the megakaryocyte-erythroid-mast cell compartment significantly expanded in the fetal liver at the expense of the B cell lineage (**Figure 1D**). Other non-disomic karyotypes also perturbed the composition of the haematopoietic compartment, albeit subtly and in different directions. An exception was the triploid fetal liver, including triplication of chromosome 21, which exhibited similar (also statistically significant) perturbations to those observed in trisomy 21. This suggests that the expansion of the megakaryocyte-erythroid-mast cell compartment observed in the triploid fetal liver is predominantly driven by trisomy 21, consistent with a view that the addition of chromosome 21 directly underpins perturbed haematopoiesis. To further evaluate the transcriptomic impact of trisomy 21 relative to other karyotypes, we compared gene expression between non-disomic cells and their diploid counterparts. Across all abnormal karyotypes and cell types, we observed genome-wide transcriptional perturbation, with most genes on the trisomic chromosomes exhibiting increased expression (**Supplementary Figure 2A, Supplementary Table 2**). Trisomy 21 had a less pronounced impact on global transcription compared to other single trisomies (**Figure 1E**). Nonetheless, the transcriptional changes in trisomy 21 cells were largely karyotype-specific (**Figure 1F**). Overall, these analyses demonstrate that the fetal haematopoietic compartment is sensitive to numerical variations in the constitutional chromosome complement, with specific effects imparted by trisomy 21.

### Stepwise transcriptional progression towards ML-DS

We set out to characterise transcriptional changes at each step along the development of ML-DS, building on single transcriptomes we generated from nine children with TAM and 11 with ML-DS (**Figure 2A**, **Table 1, Supplementary Table 1**). In the first instance, we performed Louvain clustering of all cells together and generated a Uniform Manifold Approximation and Projection (UMAP) representation of 375,548 high quality cells across the dataset (**Figure 2B, Supplementary Figure 3**). We were able to unambiguously identify clusters of neoplastic TAM and ML-DS by interrogating the nucleotide sequences derived from scRNA-seq for the presence of *GATA1* mutations (**Figure 2C, Supplementary Figure 4A**). As is usually seen in UMAPs of single neoplastic cell transcriptomes, neoplastic cells mostly formed patient specific clusters (**Supplementary Figure 3B**), further indicating that TAM blasts are transcriptionally transformed.

**Figure 2:**
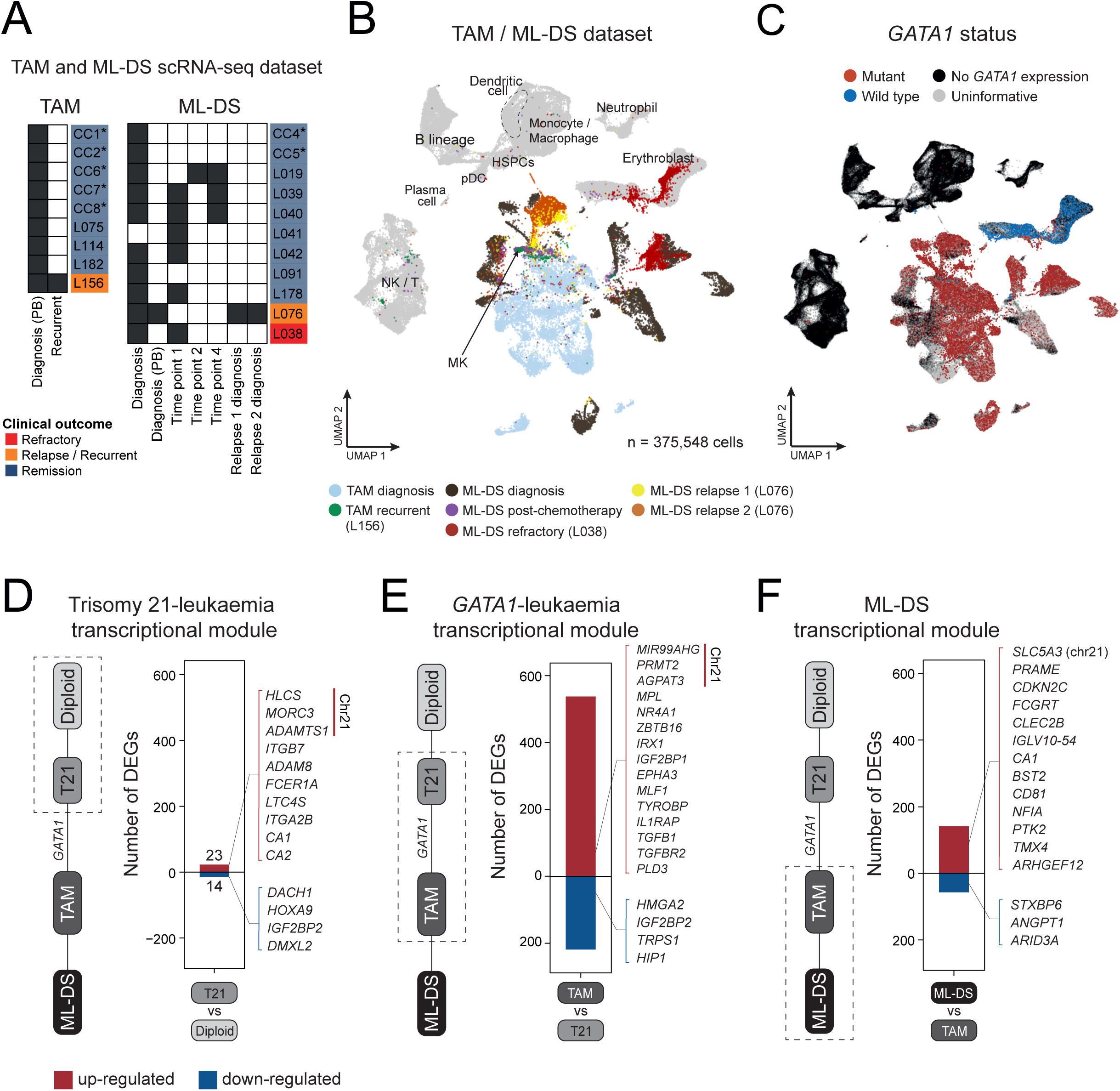
Stepwise transcriptional progression towards ML-DS. (A) TAM and ML-DS samples collected for scRNA-seq, either from peripheral blood (PB) or bone marrow aspirates at diagnosis or various timepoints post-chemotherapy. Asterisks (*) indicate FACS-sorted samples to retain only leukaemic blasts for sequencing. (B) UMAP visualisation of the TAM / ML-DS scRNA-seq dataset, with cells (dots) coloured by broad cell-type categories. Non-leukaemic cells are in grey, with the labels indicating their categories. HSPCs - haematopoietic stem and progenitor cells; MK - megakaryocyte; NK / T - natural killer cell / T cell; pDC - plasmacytoid dendritic cell. (C) UMAP visualisation of the TAM / ML-DS scRNA-seq dataset, highlighting cells with the *GATA1* mutation (red dots) all within the leukaemic blast clusters. (D) Bar plot showing number of genes in the trisomy 21 - leukaemia transcriptional module, defined as differentially expressed genes (DEGs) between trisomy 21 and diploid fetal liver MEMP/MEP contained within the ML-DS transcriptome (which are DEGs between diagnostic ML-DS cancer cells and diploid fetal liver MEMP/MEP, see **Methods**). Red indicates up-regulated genes and blue indicates down-regulated genes in trisomy 21 fetal liver MEMP/MEP (and ML-DS blasts) compared to diploid fetal liver MEMP/MEP. Highlighted are selected up- and down-regulated genes from the module with known relevant functions reported in the literature. The full gene module is detailed in **Supplementary Table 3**. (E) Bar plot showing number of genes in the *GATA1* - leukaemia transcriptional module, representing the contribution of *GATA1* mutations towards ML-DS transcriptome. The module is defined as differentially expressed genes (DEGs) between TAM and trisomy 21 bone marrow MEP contained within the ML-DS transcriptome (which are DEGs between diagnostic ML-DS blasts and trisomy 21 bone marrow MEP, see **Methods**). Red indicates up-regulated genes and blue indicates down-regulated genes in TAM blasts (and ML-DS blasts) compared to trisomy 21 bone marrow MEP. Highlighted are selected up- and down-regulated genes from the module with known relevant functions reported in the literature. The full gene module is detailed in **Supplementary Table 4**. (F) Bar plot showing number of genes in the ML-DS transcriptional module, defined as differentially expressed genes between all diagnostic (treatment naive) ML-DS blasts against conventional (i.e. non-recurrent) TAM blasts (see **Methods**). Red indicates up-regulated genes and blue indicates down-regulated genes in ML-DS blasts compared to TAM blasts. Highlighted are selected up- and down-regulated genes from the module with known relevant functions reported in the literature. The full gene module is detailed in **Supplementary Table 5**.

Next, we performed a series of differential gene expression analyses to quantitatively determine the transcriptional changes imparted by trisomy 21 followed by *GATA1* mutations along the progression towards ML-DS (see **Methods**). We first defined the ML-DS transcriptome as the difference between ML-DS cancer cells and diploid fetal liver megakaryocyte-erythroid-mast progenitor cells (MEMP/MEP). These differentially expressed genes represent the transcriptional changes between a completely normal diploid state and leukaemia. We found that only 37 genes (23 up-, 14 down-regulated), accounting for 12% of the ML-DS transcriptome, were significantly perturbed in the same directions in trisomy 21 fetal hepatic MEMP/MEP cells when compared to the diploid state (**Figure 2D, Supplementary Table 3**). Among those upregulated were five genes on chromosome 21, along with several other genes previously implicated in leukaemogenesis through promoting cell proliferation (*ITGB7*^24^, *ADAM8*^25,26^), as well as genes known to be highly expressed in mature megakaryocytes (*ITGA2B*^27^), mast cells (*FCER1A*^28^*, LTC4S*^29^) and erythroblasts (*CA1*, *CA2*^30^). Additionally, some haematopoietic progenitor markers such as *HOXA9*^31^ and *DACH1*^32^ were downregulated, which may suggest a more differentiated transcriptional state associated with ML-DS blasts.

We then assessed the contribution of *GATA1* mutation towards the leukaemic transcriptome. This is defined as the transcriptional changes between trisomy 21 bone marrow MEP and ML-DS blasts which have already occurred in TAM upon *GATA1* mutation acquisition. The analysis revealed that the majority (>80%) of differentially expressed genes between ML-DS blasts and trisomy 21 MEP are in fact already significantly perturbed in TAM blasts (**Figure 2E, Supplementary Table 4**). Eight genes on chromosome 21 were found to be significantly upregulated in cells with *GATA1* mutation, including those previously described to be involved in ML-DS (*MIR99AHG*^33^, host gene of miR-125b-2) or general cancer development (*PRMT2*^34^*, AGPAT3*^35^). Moreover, *GATA1* mutation resulted in up-regulation of its downstream targets (IRX1^36^), along with many genes normally expressed in definitive megakaryocytes, mast cells (*MPL, NR4A1*) or myeloid lineage (*ZBTB16*), but down-regulation of erythroid-specific genes (**Supplementary Figure 5A**). This is consistent with data on functional consequences of *GATA1* mutation from *in vitro* assays of patient-derived iPSCs^37^ and mouse models^33,38,39^. All three lines of evidence demonstrate that *GATA1* results in impaired erythropoiesis while promoting a myelo-megakaryocytic bias. Many genes involved in stem cell maintenance and leukaemogenesis were also up-regulated (*IGF2BP1, EPHA3, MLF1, IL1RAP, TGFB1, TGFBR2*).

Nonetheless, trisomy 21 and *GATA1* mutations together did not fully account for the ML-DS transcriptome. To distil the remaining differences required for the final step of leukaemogenesis, we compared transcriptional profiles of diagnostic (treatment naive) ML-DS cancer cells to non-recurrent TAM. We identified 198 genes differentially expressed across multiple ML-DS and TAM individuals, many of which have known roles in myeloid leukaemia, such as *ARID3A*^40^ – the downstream target of miR-125b (**Figure 2F, Supplementary Table 5**). However, we observed large patient-specific transcriptional changes when comparing blasts from each ML-DS individual against TAM (**Supplementary Figure 5B)**. These observations indicate that *GATA1* mutations explain most of the ML-DS transcriptome, yet a final transcriptional change is required to complete the evolution of ML-DS.

### Specificity of the Trisomy 21 -TAM - ML-DS transcriptional progression

Having delineated stepwise transcriptional changes along the progression of ML-DS, we set out to assess whether these signals are specific to ML-DS or shared amongst other leukaemias. To address this, we generated and examined a carefully curated dataset of 19 relevant leukaemias (159,217 single-cell transcriptomes) for comparison with TAM and ML-DS (as detailed in **Table 1**, **Figure 3A, Supplementary Figure 6A**), including: rare cases of diploid myeloid neoplasm with and without *GATA1* mutation (**Supplementary Figure 6B-C**), which enabled us to disentangle transcriptional effects of trisomy 21 and *GATA1*; morphologically and cell type related leukaemias (acute megakaryocytic leukaemia - AMKL, acute erythroid leukaemia - AEL, other myeloid leukaemias); as well as lymphoid neoplasms with and without trisomy 21, to tease apart transcriptional effects of lineage (lymphoid versus myeloid) and trisomy 21. In addition, we included in our analyses relevant normal cell types from independent (published) datasets of fetal liver (diploid)^11^ and fetal bone marrow (both diploid and trisomy 21)^12^ cells. Finally, we generated an orthogonal bulk transcriptomic dataset for further validation, consisting of TAM (n=16), ML-DS (n=13), other myeloid leukaemias (n=38), along with flow-cytometry-sorted normal diploid haematopoietic progenitors (**Table 1, Supplementary Table 1**).

**Figure 3:**
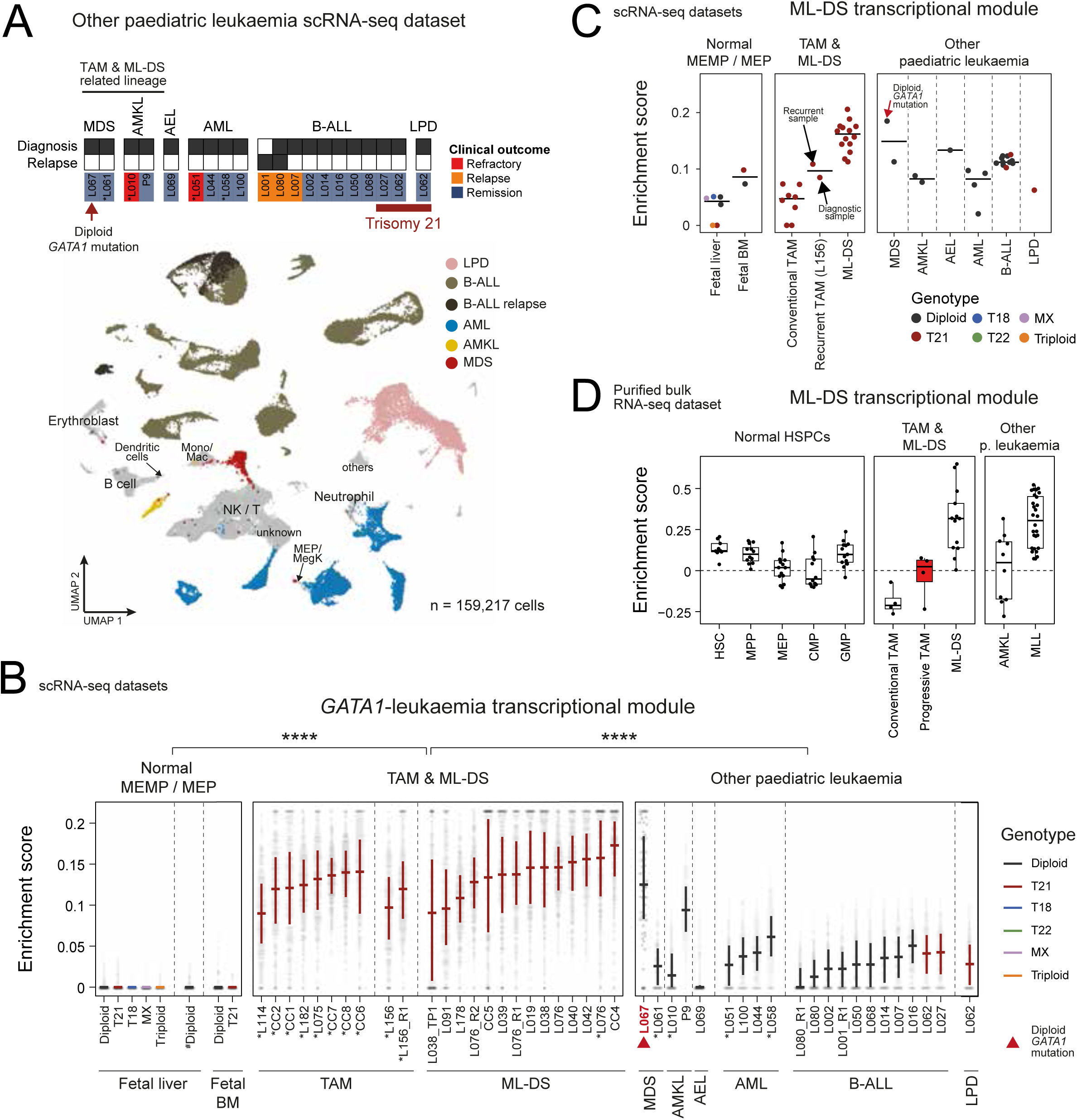
Specificity of the Trisomy 21 - TAM - ML-DS transcriptional progression. (A) Top - Other paediatric leukaemia samples collected for scRNA-seq, obtained either at diagnosis or relapse diagnosis (denoted as “Relapse”). Asterisks (*) indicate peripheral blood samples; all others are bone marrow aspirates. Bottom - UMAP visualisation of the dataset, with cells (dots) coloured by broad cell-type categories. Non-leukaemic cells are in grey, with the labels indicating their categories. (B) Enrichment score of the *GATA1* - leukaemia transcriptional module (y-axis) across individual cells (grey dots) from different scRNA-seq datasets (x-axis) consisting of: normal MEMP/MEP with different karyotypes from the fetal liver (including diploid cells from Popescu, D-M. *et al.*, 2019^11^, denoted by #) and fetal bone marrow (BM) from Jardine, L. *et al*., 2021^12^; TAM/ML-DS blasts; and leukaemic blasts from other leukaemias. Red arrow highlights diploid MDS sample with *GATA1* mutation. Leukaemic cells are grouped by patient, timepoint (initial diagnosis unless indicated otherwise in the group name) and tissue. Amongst leukaemia samples, asterisks (*) indicate peripheral blood samples; all others are bone marrow aspirates. Cross bars indicate the interquartile range of enrichment score distribution, coloured by corresponding donors’ germline genotypes. The mean enrichment score per individual across TAM/ML-DS cohort is significantly higher than that in normal MEMP/MEP group (p = 9.507e-08, one-sided Wilcoxon rank sum test), as with AML/B-ALL group (p = 9.884e-11, one-sided Wilcoxon rank sum test). (C) Per-sample median enrichment score of the ML-DS transcriptional module (y-axis) across different scRNA-seq datasets as detailed in (B) (x-axis). Dots represent the median module score across normal MEMP/MEP cells with different karyotypes from fetal liver and fetal bone marrow (BM), or blasts from each patient, coloured by corresponding donors’ germline genotypes. Black horizontal bars represent group-level medians. The black arrow highlights a recurrent TAM case, where the median enrichment score is distinctly higher compared to other TAM cases. Red arrow highlights diploid MDS sample with *GATA1* mutation. (D) Per-sample enrichment score of the ML-DS transcriptional module (y-axis) in an independent bulk RNA-seq dataset composed of FACS-sorted normal haematopoietic stem and progenitor cells (HSPCs), TAM/ML-DS diagnostic samples, and leukaemic blasts from other paediatric leukaemias. ***Abbreviation*** <u>Leukaemia</u>: MDS - myelodysplastic syndrome; AMKL - acute megakaryoblastic leukaemia; AML - acute myeloid leukemia; AEL - acute erythroid leukemia; B-ALL - B cell acute lymphoblastic leukaemia; LPD - lymphoproliferative disorder; MLL - mixed lineage leukaemia. <u>Sample timepoint</u>: R - recurrent; R1 - relapse 1 diagnosis; R2D - relapse 2 diagnosis; TP1 - timepoint 1, TP2 - timepoint 2, TP4 - timepoint 4. <u>Cell types</u>: HSPCs - haematopoietic stem and progenitor cells; HSC - haematopoietic stem cell; MPP - multipotent progenitor; MEP - megakaryocyte-erythroid progenitor; CMP - common myeloid progenitor; GMP - granulocyte-monocyte progenitor; NK / T - natural killer cell / T cell. <u>Genotypes</u>: T - trisomy; MX - monosomy X.

We calculated the enrichment score for each gene set of transcriptional changes we had defined to delineate the evolution of ML-DS from disomic liver cells. First, we examined imprints of the trisomy 21 transcription module (as defined in **Figure 2D**). Amongst normal fetal MEP cells, these imprints were specific to trisomy 21. In neoplasms we found an enrichment in leukaemias of the MEP lineage (MDS with *GATA1* mutation, AMKL, AEL; **Supplementary Figure 7A**). By contrast, strong enrichment of the *GATA1* transcription module (as defined in **Figure 2E**) was confined to diploid MDS with *GATA1* mutation, indicating that the *GATA1* module does not depend on trisomy 21 (**Figure 3B**). Of note, AMKL cancer cells exhibited an intermediate signal, which is most likely driven by the megakaryocytic genes upregulated as a result of the *GATA1* mutation (**Supplementary Figure 5A**). Assessment in the independent bulk RNA-seq dataset validated these observations (**Supplementary Figure 7B-C**).

Finally, assessing the ML-DS module (as defined in **Figure 2F**), we found an enrichment across all leukaemias, including those from the B-lineage (**Figure 3C, Supplementary Figure 8A**). This suggests that the ML-DS module may represent a gene set widely dysregulated in leukaemia that is relevant beyond the transformation of TAM to ML-DS. Consistent with this notion, we found established leukaemogenic genes such as *CD81*^41^, *BST2*^19,42^ or *PTK2*^43^ contained in the ML-DS module (**Supplementary Figure 8B**, **Supplementary Table 5**).

Of note, we also observed imprints of this cancer transcriptome in a child with “recurrent TAM” which, from a clinical treatment perspective, probably represents ML-DS rather than TAM (black arrow in **Figure 3C**). Extending this observation into the independent bulk cohort, where 4 TAM cases eventually progressed to ML-DS (termed “progressive TAM”), we observed the same pattern (**Figure 3D**) that the ML-DS transcriptomes may delineate progressive TAM.

### Genetic and transcriptional evolution of progressive ML-DS

In a final analysis, we examined rare cases of progressive, ultimately fatal ML-DS: one child (L076) with two relapses and one child (L038) who developed refractory disease following the first course of chemotherapy. With samples from all timepoints at our disposal, we were able to examine both the genetic and transcriptional evolution of progressive ML-DS. We determined genetic evolution through whole genome sequencing based phylogenies using established analytical approaches^44^. We matched genetic clones with single cell mRNA clusters by assessing allelic imbalances of cancer-defining copy number changes in mRNA reads, which is a precise and expression-independent approach for identifying cancer cells in single cell mRNA sequencing data^45^.

Following diagnosis with ML-DS, child L076 had two relapses after short lived remissions (months) on each occasion. Phylogenetic analyses showed that the two relapses were derived from the diagnostic clone via a common precursor but then diverged in their genetic development, accruing mutations in parallel (**Figure 4A, Supplementary Table 6**). The copy number profiles of the major clones of all three timepoints were identical (**Figure 4B, Supplementary Figure 9A-B**), and none of the additional substitutions or indels generated plausible driver events. Interestingly, the second relapse exhibited a mutational signature of prior treatment exposure (**Figure 4A**). Consistent with the genetic distinction between initial disease and relapses, transcriptionally both relapses segregated clearly from the diagnostic clone (**Figure 4B**). However, despite their subsequent parallel evolution, they remained transcriptionally similar to each other. This may indicate that these parallel clones retained a common transcriptional phenotype of progressive disease, or that both lineages converged on a common phenotype. Examining differential expression analyses between progressive disease and the initial diagnostic clone, revealed a range of expression changes that have been implicated in cancer cell survival or treatment resistance such as upregulation of *FHL2*^46^*, CXCL8*^47^*, EPS8*^48^ and *CD82*^49,50^ (**Figure 4C, Supplementary Table 8**). Overall, therefore, progressive disease in this child (L076) segregated genetically and transcriptionally from the initial diagnostic clone, which was underpinned by plausible expression changes without any specific mutation accounting for aggressive disease. By contrast, in child L038 who developed refractory disease after one course of chemotherapy, we identified a plausible progression event, namely deletion of *TP53* (through chromosome 17p deletion) which has previously been implicated as a driver of aggressive ML-DS^21^ (**Figure 4D, Supplementary Figure 9C).** We found that progressive disease evolved from a clone with *TP53* deletion that had been present at diagnosis (**Figure 4E, Supplementary Figure 9D, Supplementary Table 7**). However, this refractory clone had not been derived from the diagnostic clone but represented a parallel lineage. The transcriptional difference between ML-DS cells with, or without deletion of *TP53*, revealed a number of expression changes that plausibly contributed to disease progression, including some in common with expression changes of progression in the first child, L076 (**Figure 4C, Supplementary Table 8**). We observed specific down-regulation of genes such as *LDLR* and *CD81* in relapse/refractory blasts compared to the initial diagnostic clones. Importantly, genes underpinning these changes did not reside on the deleted chromosome 17p, although we were able to clearly observe transcriptional imprints of 17p deletion in refractory blasts (**Supplementary Figure 9E**). Finally, we examined transcriptional commonalities between blasts from all samples of these two children which showed that, irrespective of disease stage, blast transcriptomes retained *GATA1* induced gene expression (**Figure 2E**), underscoring our proposition that they may represent a common vulnerability across the ML-DS disease spectrum.

**Figure 4:**
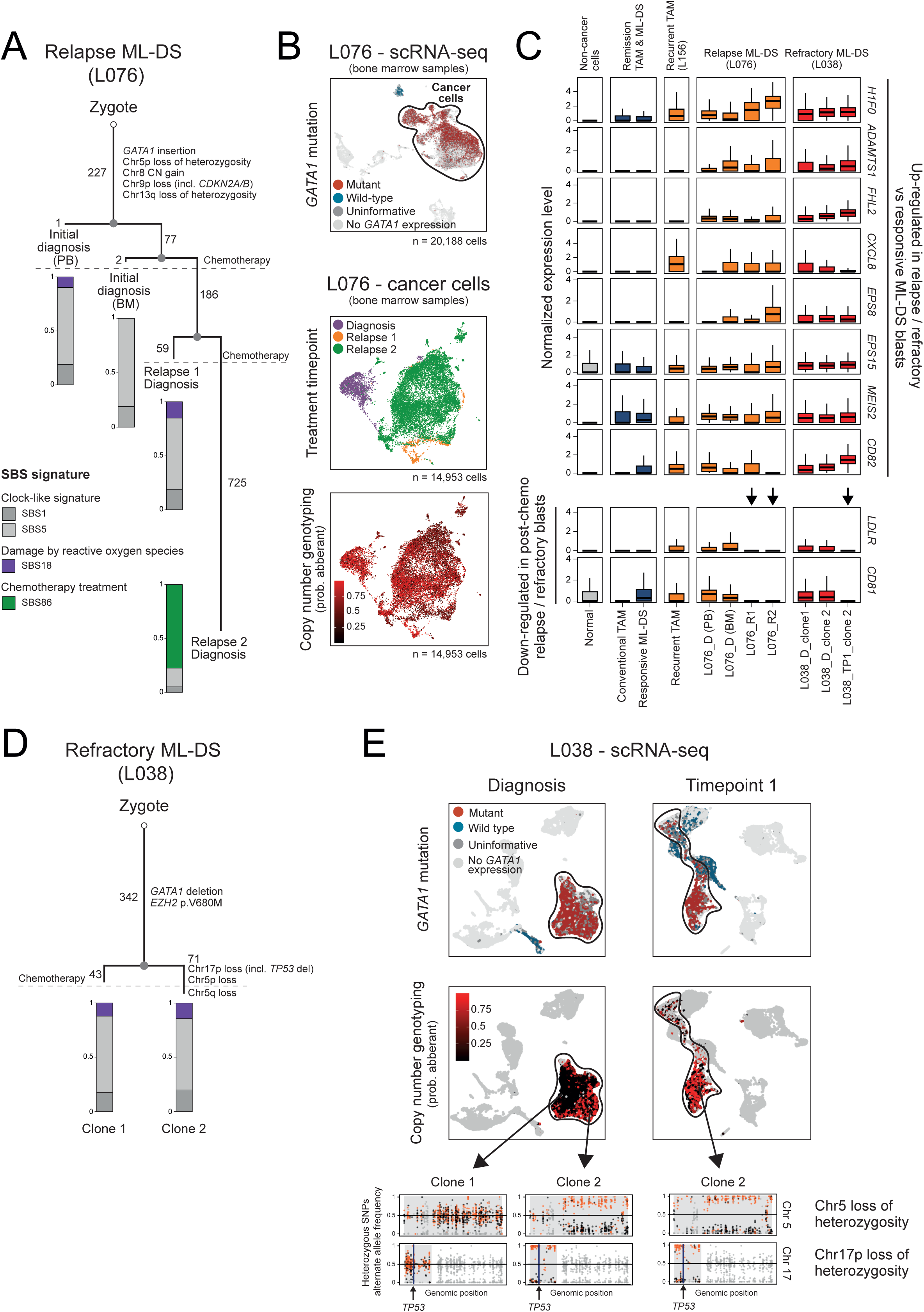
Genetic and transcriptional evolution of progressive ML-DS. (A) Reconstructed phylogeny of clones in child L076 with two relapses, annotated with known driver mutations, copy number variants, and estimated number of single-nucleotide substitutions. Bar plots show the proportion of single-base substitution signatures. (B) UMAP visualisation of scRNA-seq data of bone marrow samples from child L076. All single transcriptomes (dots) obtained from bone marrow biopsies at diagnosis, first and second relapse diagnosis are shown. Cells are coloured by *GATA1* mutation status (top), sample collection timepoint (middle), and probability of having copy number aberrations (bottom). (C) Assessment of marker genes in relapse and/or refractory ML-DS blasts (post-chemotherapy) compared to treatment-naive ML-DS blasts, either from the same cases (L076 and L038) or from other cases that achieved remission following chemotherapy. Box plots show the distribution of single-cell normalised expression (y-axis) for selected markers (rows) previously implicated in cancer cell maintenance or treatment resistance. Cells are grouped by identity (normal or cancer) and the corresponding donor’s treatment outcome (x-axis). (D) Reconstructed phylogeny of clones in child L038 with treatment-refractory disease. The phylogeny is annotated with known driver mutations, copy number variants, and the estimated number of single-nucleotide substitutions. Bar plots show the proportion of single-base substitution signatures, with the same colour scheme as in (A). (E) UMAP visualisation of scRNA-seq data of patient L038. All single transcriptomes (dots) captured from both timepoints are shown. Cells are coloured by *GATA1* mutation status (top) and the probability of having copy number aberrations (middle). Bottom panel shows the aggregated B-allele frequency of heterozygous single-nucleotide polymorphisms (SNPs) across different clones from each timepoint. Each dot represents a heterozygous SNPs along chromosomes 5 and 17 (x-axis), and the corresponding aggregated B-allele frequency (y-axis) are shown. Orange dots indicate alternate alleles located on the major chromosome, whereas black dots indicate alternate alleles located on the minor chromosome. Copy number segments are grey-shaded regions. ***Abbreviation*** Chr – chromosome <u>Sample timepoint</u>: R - recurrent; R1 - relapse 1 diagnosis; R2D - relapse 2 diagnosis; TP1 - timepoint 1. <u>Tissue</u>: PB - peripheral blood; BM - bone marrow.

## DISCUSSION

We studied the genetic and transcriptional evolution of ML-DS directly from natural variation which enabled us to delineate transcriptional effects of individual genetic steps and measure their contribution to the ML-DS transcriptome. A key finding of our analyses was that *GATA1* induced transcriptional changes predominated the entire disease spectrum, including relapse/refractory disease. The genetic changes that underpin the progression from TAM to ML-DS do not override or erase the *GATA1* transcriptome. This indicates that *GATA1* mediated expression remains relevant at every disease stage and may thus represent a persistent therapeutic vulnerability.

The strength of our work lies in leveraging natural variation from primary disease samples as the basis of our investigation. High throughput single cell mRNA sequencing enabled us to extract relevant cell populations for precise expression analyses to distil transcriptional effects of genetic variation. We studied several individuals with TAM and ML-DS, including cases of progressive ML-DS, which enabled us to derive patient-overarching effects, which we further validated in an independent bulk transcriptomic dataset. A key limitation of our approach is its dependence on natural variation. Accordingly, some of the comparators in our analyses are individual cases of conditions that are vanishingly rare, such as a fetus with whole genome triploidy or a disomic child with *GATA1* driven MDS. However, even in these cases, single cell transcriptomics delivered multiple readouts, which substantiates our findings at the level of individual patients.

Previous work has explored the functional significance of each genetic aberration along the development of ML-DS through *in vitro* assays using primary human haematopoietic stem and progenitor cells, mouse models or xenotransplantation in mice^1,33,39,40,51–54^. Here, we examined the contribution of each genetic step in the transcriptional evolution of ML-DS, and showed that *GATA1* mutations account for most of the ML-DS transcriptome. This held true in progressive ML-DS in two children. Despite a complex phylogenetic relationship between cancer clones and transcriptional changes along each step, the *GATA1* transcriptome continued to predominate in ML-DS-blasts. *GATA1* induced changes were independent of constitutional trisomy 21 and specific to *GATA1* mutated neoplasms, supporting previous reports that trisomy 21 is not essential for leukaemogenesis^33,55^. This observation would explain why ML-DS can only be generated via *GATA1* mutations, as well as clarifying rare cases of TAM/ML-DS like neoplasms with *GATA1* mutation in children without Down syndrome^55^. Unlike the highly disease specific effects of *GATA1* mutation, ML-DS defining transcriptional changes represented a generic leukaemogenic transcriptome, underscoring the fact that ML-DS is a cancer. Given the central contribution of *GATA1* mutation associated transcriptional changes to ML-DS, it is conceivable that therapeutic interference with these changes may disintegrate, and thus treat, the ML-DS transcriptome.

Owing to meticulous tissue banking efforts, we were able to study rare clinical phenotypes of progressive TAM and ML-DS. Recurrent TAM is a largely cryptic disease which clinically is indistinguishable from ML-DS and often, on clinical grounds, will eventually be treated as ML-DS. No molecular differences between progressive and non-progressive TAM have been described to date, and there are no effective risk stratification strategies^56–59^. Remarkably, both single-cell transcriptomes and bulk transcriptomes of progressive TAM were transcriptionally different from conventional (i.e. non-recurrent) TAM and more closely resembled ML-DS. This indicates that we may have identified potential biomarkers that enable the prospective identification of progressive TAM.

Our quantitative analysis in primary human samples, integrated with DNA analyses, details a unified landscape of the transcriptional evolution underpinning ML-DS. Some of our findings, such as transcriptional features of progressive TAM or the pre-existence of a refractory clone at diagnosis in one child, may be of clinical utility. Pursuing these clinical hypotheses further will require large scale, international prospective investigations. At the same time, our work outlines a broadly applicable analytical approach for distilling transcriptional effects of individual mutations directly from human cancer cells and natural variation rather than model systems.

### Resource availability

WGS and single-cell mRNA sequencing data has been deposited in the European Genome-Phenome Archive (EGA) under the accession codes EGAD00001015453 (WGS) and EGAD00001015452 (scRNA-seq). Processed counts for all single-cell and bulk transcriptomic data are available in the Gene Expression Omnibus (GEO) repository under the accession code YYYY*. Raw sequencing data of the bulk transcriptomic dataset is available upon request from Prof. Jan-Henning Klusmann

*(* all accession codes will be provided prior to publication)*

All code used to reproduce the analysis and figures described in this manuscript is available at https://github.com/miktrinh/ML-DS.

## Acknowledgements

This study was funded by the Wellcome Trust (institutional grant to the Wellcome Sanger Institute (WT206194), and personal fellowship to S.B. (223135/Z/21/Z)). This study was also supported by “Hilfe für krebskranke Kinder Frankfurt e.V.” as part of the C^3^OMBAT-AML consortium, the German Research Foundation (DFG, KL 2374/8-1) and the European Research Council (ERC) under the European Union’s Horizon 2020 Research and Innovation Programme (grant agreement No. 714226). Additional funding was received from the Wenner-Gren Foundations (personal fellowship to A.W.). We are indebted to the children and families who participated in this study.

## Author contributions

S.B., J.B., and J.H.K. conceived and co-directed the study. M.K.T. performed overall bioinformatic data analysis and interpreted the results, aided by M.Y., L.J., K.S., A.W., N.D.A., H.J.W., T.D.T, A.H., K.S., J.G.D., S.A. and D.O.C.. H.I., R.T., C.P., A.O., T.O., D.Z., L.M., E.P., E.R., A.H., K.S., and S.A. collected and processed the samples for sequencing experiments. S.B. and M.K.T. wrote the manuscript. All authors have read and approved the manuscript.

## METHODS

### Sample acquisition, ethics and patient consent

Fetal liver samples were obtained from the MRC–Wellcome Trust - funded Human Developmental Biology Resource (HDBR; http://www.hdbr.org^60^) with appropriate written consent and approval from the North East – Newcastle & North Tyneside 1 Research Ethics Committee. HDBR is regulated by the UK Human Tissue Authority (HTA; www.hta.gov.uk) and operates in accordance with the relevant HTA Codes of Practice.

Leukaemia samples from bone marrow aspirates or peripheral blood were collected through the Great Ormond Street Hospital for Children diagnostic archives, as approved by a National Research Ethics Service Committee (London Brent, reference 16/LO/0960); and the AML Berlin-Frankfurt-Münster (BFM) study group. Bulk transcriptomic processed data was obtained with approval from the Goethe University Frankfurt (Ethics #2021-341 and #2023-1565). Ethical approval was obtained from the London - Brent Research Ethics Committee, the University Duisburg-Essen (Ethics #17-7462), Hannover Medical School (Ethics #6741M) and Martin-Luther-University Halle-Wittenberg (Ethics #2019-103). Relevant clinical information of the patient cohort is included in **Supplementary Table 1**. Informed consent to participate in research was obtained from all patients or their legal guardians as stipulated by the study protocols.

### TAM / ML-DS blast enrichment by flow cytometry

TAM and ML-DS samples obtained from the AML Berlin-Frankfurt-Münster (BFM) study group were FACS-sorted to obtain pure leukaemic blasts prior to the 10X workflow (**Supplementary Table 1**). Mononuclear cells were enriched with density gradient centrifugation and subsequently FACS sorted. Blast was defined as CD45dim/CD33+/CD117+/CD7+/CD11a-.

### 10X single cell RNA sequencing (scRNA-seq)

Fresh fetal livers were dissociated into single-cell suspensions. For the leukaemia samples (bone marrow aspirates or peripheral blood), peripheral blood mononuclear cells were prepared by density centrifugation using Lymphoprep (Stemcell) according to manufacturer’s instructions, also generating single-cell suspensions. The suspension was passed through a 70μm cell strainer (Falcon) and washed with PBS. If necessary, live cell enrichment using a Dead Cell Removal kit (Miltenyi Biotec) and red blood cell removal using the eBioscience 10X RBC Lysis Buffer (Multi-species) were performed as per manufacturer’s instructions. For some fetal liver samples, cells are further sorted into CD45+ and CD45-fractions using magnetic beads (detailed in **Supplementary Table 1**). All enriched live cells were washed and counted using a hemocytometer with trypan blue, single cell suspensions were adjusted to 1000 cells/ul accordingly. Cells were loaded onto the Chromium 10X controller as per the standard protocol of the Chromium Single Cell 3’ Reagent kits (V3 chemistry) or 5’ Reagent kits (v2 chemistry) in order to capture between 7000 cells/chip position (detailed in **Supplementary Table 1**). All the following steps were performed according to the standard manufacturer protocol. Post GEM-RT clean-up, cDNA amplification, and 5′ gene expression library construction were carried out according to the manufacturer’s instructions. The resulting libraries were sequenced on the Illumina Novaseq 6000 platform, aiming for an average of 300,000 reads per cell.

### Quality control and preprocessing of scRNA-seq data

Raw sequencing data was processed and mapped to the reference genome (GRCh38 1.2.0 or GRCh38 2020-A as detailed in **Supplementary Table 1**), using Cell Ranger pipeline^61^. The filtered count matrix, outputted by Cell Ranger, was then further QC’ed using Seurat^62^ (v4.0.1) in R (v4.0.4). Cells with <300 genes, <500 UMIs, or mitochondrial fraction exceeding 30% were removed. Scrublet^63^ (v0.2.3) was used to identify doublets. Cells were excluded if identified as doublets by Scrublet, or having a doublet score > 0.5. Ambient mRNA contamination was removed with SoupX^64^ (v1.6.1). High resolution clusters (resolution=10) with >50% cells failing QC were also excluded.

The data were log normalised and scaled, and principal components were calculated using highly variable genes, following the standard Seurat workflow. Louvain clustering was performed (resolution = 1), and a uniform manifold approximation and projection (UMAP) calculated, using the top 75 principal components. No integration or batch correction method was performed to preserve the biological variation across different karyotypes and cancer samples.

### Bulk mRNA sequencing and processing

RNA was isolated from cells using the Quick-RNA Miniprep Kit (Zymo Research). Paired-end libraries with 2 × 50, 75, 100 and 150 bp reads were prepared from the extracted RNA using the TruSeq Stranded total RNA LT Sample Prep (RiboZero Gold, Illumina) using Illumina Methodology by Novogene Company, Ltd. Raw sequencing reads were trimmed with fastp^65^ (v0.24.0) and aligned to GRCh38 using STAR^66^ (v2.6.0). Mapped reads were annotated using Ensembl v.108. Gene expression levels in transcripts per million (TPM) were quantified from the mapped reads using salmon^67^(v1.10.3).

### Whole genome sequencing

Whole genome sequencing (WGS) was performed on the fetal tissues with abnormal karyotypes, and those from two cases of aggressive ML-DS: one relapsed twice, and one refracted to standard chemotherapy (details in **Supplementary Table 1**). Bulk DNA sequencing was performed as previously described^8^. In brief, DNA was extracted using the AllPrep DNA/RNA/Protein Mini Kit (QIAGEN) following the standard protocol, and short insert (∼500bp) genomic libraries were generated. Finally, 150 bp paired-end sequencing was conducted on the Illumina NovaSeq 6000 platform according to Illumina’s standard library generation protocols (with PCR). The average sequence coverage was at least 30X per sample (details in **Supplementary Table 1**).

Raw DNA sequencing data were aligned to the GRCh38 (Ensembl 103) reference genome using the Burrows-Wheeler algorithm (BWA-MEM)^68^.

### Sample-level genotyping

To ensure that sequencing data (both WGS and scRNA-seq) from the same individuals are correctly labelled, sample-level genotyping was performed using the *matchBAMs* function from the R package *alleleIntegrator*^45^.

The karyotype of each donor was also confirmed. For those with WGS data (see **Supplementary Table 1**), we compared the coverage of affected chromosome(s) to the rest of the genome as follows:

- Single-chromosome trisomy: coverage of the trisomic chromosome is approximately 1.5 times higher than other disomic chromosomes (other copy number regions were excluded, if any).
- Monosomy X: There is no coverage on chromosome Y (confirming that the individual is a female), and the coverage of chromosome X is half that of other disomic chromosomes.
- Whole genome trisomy: We confirmed that the coverage is higher than expected (submitted 100X but got 130X?). We further validated this by assessing the distribution of the alternate-allele frequency at heterozygous single nucleotide polymorphisms (SNPs), which showed a bimodal distribution with peaks around 1/3 and 2/3.

For cases with only scRNA-seq data, we first combined all scRNA-seq samples from the same individual, and identified the individual-specific heterozygous SNPs using the R package *alleleIntegrator*^45^. We then calculated the alternate/B-allele frequency (BAF) at heterozygous SNPs, aggregated across all high-quality sequencing reads (mapping quality >= 200, base quality >= 20, minVarQual = 225) on each chromosome. By assessing BAF distributions for each individual, we confirmed that trisomic chromosomes exhibit a bimodal BAF distribution with peaks around 1/3 and 2/3, while disomic chromosomes show a unimodal distribution around 0.5.

### Cell type annotation

#### Fetal liver scRNA-seq dataset

A semi-automated cell type annotation using a label transfer approach was employed. Using Celltypist^69^, a logistic regression model was trained on the reference scRNA-seq atlas of the human fetal livers (Popescu, DM., Botting, R.A., Stephenson, E. *et al.*, 2019^11^). The model was then used to calculate predicted similarity scores for individual cells in our query fetal liver dataset against each reference class (i.e. different cell types in the reference dataset) (**Supplementary Figure 1D**). Cells were assigned the label of the class with the highest positive similarity score. Annotation was further refined by manually assessing the expression of well-established cell-type specific marker genes (**Supplementary Figure 1C**) in high-resolution Louvain clusters.

To quantify the proportion of different haematopoietic lineages in the fetal livers of different karyotypes, we first excluded all non-haematopoietic cells, including: endothelial cells, fibroblasts, hepatocytes, cholangiocytes, mesothelial cells, neurons, and trophoblasts. The remaining cells were grouped by donor ID and corresponding lineages as follows:

- HSC/MPP
- Megakaryocyte – Erythroid – Mast cell lineage: MEMP/MEP, early megakaryocytes, megakaryocytes, early/mid/late erythroblasts, mast cells.
- B lineage: LMPP/ELP, pro B cells, pre B cells, B cells.
- Myeloid lineage: CMP/GMP, DC1, DC2, pDC, pro-monocytes, monocytes, macrophages, Kupffer cells, pro-myelocytes, myelocytes.
- NK/T lineage: ILC precursors, natural killer/T cells.

The proportion of each lineage in each donor was calculated as the number of cells in the lineage divided by the total number of haematopoietic cells from that donor.

To compare the lineage proportions between those with abnormal karyotypes (trisomy 21 and Triploid) and diploid fetal livers, we first calculated the respective proportions by aggregating cells across all donors within each karyotype group. We then performed a one-tailed two-sample proportion z-test, using the *prop.test* function in R, to compare proportions between diploid and each abnormal karyotype.

### Leukaemia scRNA-seq dataset

A similar approach was used to annotate both leukaemia scRNA-seq datasets (TAM/ML-DS and other leukaemias), where automated label transfer was followed by manual assessment of canonical cell type specific marker genes. However, as these datasets contain neoplastic cells that may not closely resemble normal cellular transcriptomes, we employed a logistic regression method previously detailed in Young *et al.*, 2018^70^. Specifically, a logistic regression model with elastic net regularisation (alpha = 0.99) was trained on our diploid fetal liver scRNA-seq data (reference dataset), using the *cv.glmnet* function from the *glmnet* R package. This model was applied to calculate the predicted similarity scores for individual cells in the leukaemia datasets against each reference class (i.e. different cell types in the reference dataset) (**Supplementary** Figure 3C). To allow for the possibility that some query cells may not match strongly to any of the reference cell types, softmax normalisation was not used. Cells were assigned the label of the class with the highest positive similarity score. Annotation was further refined by manually assessing the expression of well-established cell type specific marker genes (**Supplementary** Figure 3D) in high-resolution Louvain clusters.

### Identification of *GATA1* mutated neoplastic cells in TAM / ML-DS scRNA-seq data

#### Genotyping for the presence of GATA1 mutations in single-cell transcriptomes

The exact *GATA1* mutation(s) acquired in each TAM / ML-DS case was obtained from clinical history and confirmed with our WGS analyses where possible. Next, we examined the scRNA-seq reads from each sample, using samtools^71^ to identify high quality reads overlapping the *GATA1* mutation position +/- 70 bases. These reads were sorted into *GATA1* mutant or wild-type reads based on the nucleotide sequence; and assigned to their corresponding cell barcodes. The number of mutant or wild-type *GATA1* reads per cell was tallied. It is expected that only one allele of *GATA1* is expressed in a given cell, as *GATA1* is located on chromosome X outside the PAR region, combined with X-inactivation in females. Therefore, cells were classified as mutant (and therefore neoplastic cells) if they expressed only the *GATA1* mutant allele, and normal if they exclusively expressed the wild-type allele. For cells with both wild-type and mutant reads (likely due to doublets or ambient RNA contamination), cells were classified as mutant if they had at least five more mutant reads than wild-type reads, or wild-type if the reverse was true. Otherwise, they were considered ambiguous.

### Differential gene expression analysis

All differential gene expression analyses outlined below employed a donor-level pseudo-bulk approach. Donor-level pseudo-bulks were generated by aggregating raw counts per gene across all relevant cells from each donor. Pseudo-bulks with fewer than 30 cells were excluded, along with genes on chromosome Y. DESeq2^72^ was used to perform differential gene expression analysis, following the workflow outlined in the vignette [https://bioconductor.org/packages/devel/bioc/vignettes/DESeq2/inst/doc/DESeq2.html]. In brief, a negative binomial generalised linear model was fitted to estimate the effect of independent variables on gene expression levels. Wald statistics with Benjamani-Hochberg correction for multiple hypothesis testing were used to identify genes with significant differential expression (i.e. genes with log2 fold-change (log2FC) of expression level significantly different from zero). The log2FC for each gene was extracted and subjected to shrinkage correction to improve accuracy for genes with high dispersion. Genes were identified as significantly differentially expressed (DEGs) using the following criteria unless stated otherwise:

1. Adjusted p-value < 0.05
2. Absolute log2FC >= 0.5
3. Expressed in >= 10% of cells across all samples from either group

### Transcriptional consequences of different abnormal karyotypes on fetal liver cell types

To assess the transcriptional impact of different abnormal karyotypes on fetal liver cell types, we performed pseudo-bulk differential expression analyses comparing each cell type from individual abnormal karyotypes with the diploid counterpart (**Figure 1E-F; Supplementary Figure 2**). Due to low cell numbers, some cell types were grouped together:

- Megakaryocyte: early megakaryocytes and megakaryocytes.
- Myelocyte: pro-myelocytes and myelocytes.
- Monocyte: pro-monocytes and monocytes.
- B cell progenitors (“B.cell.prog”): LMPP/ELP; pro B cells; and pre B cells.
- NK/T: ILC precursors and natural killer / T cells.

To estimate the sample-specific size factor, the *estimateSizeFactors* function was used with *controlGenes* parameter specified to only genes on disomic chromosomes. For the triploid vs diploid comparison, *controlGenes* is specified as genes on chromosome X outside the PAR region (which is known to escape X-inactivation), with the assumption that despite being triplicated, the expression level of these genes is least likely to be affected compared to diploidy due to the X-inactivation mechanism in females. The model design used was “ Gene_expression ∼ Genotype + Assay “.

Genes were considered to be expressed by a given cell type if they were expressed in >= 10% diploid cells of that cell type.

### Derivation of the trisomy 21 - leukaemia transcriptional module

We defined the “trisomy 21 - leukaemia transcriptional module” as the transcriptional changes observed in trisomy 21 fetal liver MEMP/MEP cells compared to their diploid counterpart that is contained within the ML-DS transcriptome.

First, the ML-DS transcriptome is defined as DEGs between diagnostic ML-DS blasts (from bone marrow samples only) compared to diploid fetal liver MEMP/MEP cells. To minimise potential confounding artifacts, only diploid fetal liver samples sequenced using the same 10X reagent kit (Chromium Single Cell 5’ Reagent kits - v2 chemistry) as the ML-DS samples were included. The model design used was “Gene_expression ∼ Group“. We then examined which of these DEGs were already significantly perturbed in the same direction in trisomy 21 fetal liver MEMP/MEP. We performed a second pseudobulk analysis comparing expression levels of these genes in cells with trisomy 21 and diploid cells. Here, all relevant fetal liver samples sequenced with either 5’ or 3’ reagent kit were included. To account for the effects of different reagent kits, the model design used was “Gene_expression ∼ Genotype + Assay“, where “Assay” is a two-level factor representing reagent kits. Genes which met criteria (1) above were considered to be significantly differentially expressed in trisomy 21 fetal liver MEMP/MEP, and thus included in the “trisomy 21 - leukaemia transcriptional module”. Criteria (2) and (3) were excluded here as we wanted to retain all relevant changes in expression level, including very subtle perturbations represented by low log2FC values. Full results from both analyses can be found in **Supplementary Table 3**.

### Derivation of the *GATA1* - leukaemia transcriptional module

The “*GATA1* - leukaemia transcriptional module” reflects the transcriptional consequences of *GATA1* mutations observed in TAM blasts compared to trisomy 21 MEP cells, which are contained within the ML-DS transcriptome.

We first performed a pseudobulk differential expression analysis comparing diagnostic bone marrow ML-DS blasts with normal trisomy 21 MEP cells. More specifically, the latter cells were obtained from post-treatment remission bone marrow samples of four different ML-DS patients (L019, L038, L040, L041) with the highest number of captured MEP cells. DEGs were identified using all 3 criteria outlined above. We then tested for the differential expression of these genes between conventional (i.e. non-recurrent) TAM blasts and the same set of trisomy 21 MEP cells. Genes which met criteria (1) were considered to be significantly differentially expressed in TAM blasts as transcriptional changes induced by *GATA1* mutations, and thus included in the “*GATA1* - leukaemia transcriptional module”. Full results from both differential expression analyses can be found in **Supplementary Table 4**.

### Derivation of the ML-DS leukaemia transcriptional module

The “ML-DS leukaemia transcriptional module” captures the transcriptional differences between the leukaemic state of ML-DS and the pre-leukaemic state of TAM blasts. A pseudobulk differential expression analysis was performed comparing ML-DS cancer cells from diagnostic bone marrow samples against conventional TAM blasts. Genes were considered to be DEGs if all 3 criteria outlined above were met. Full results can be found in **Supplementary Table 5**.

Grouping all ML-DS cases into one category may have obscured between-patient heterogeneity, which is known to be a prominent feature across cancer types. To address this, we performed a second analysis comparing the conventional TAM group against individual ML-DS cases. DEGs for each comparison were extracted using the above 3 criteria. This analysis revealed a large number of transcriptional perturbations in ML-DS compared to TAM which are individual-specific (**Supplementary Figure 5B**).

### Characterisation of the transcriptional signature of progressive ML-DS

To investigate the transcriptional changes potentially associated with progressive ML-DS and chemotherapy resistance, we specifically looked for markers associated with chemotherapy resistance in the progressive blasts in L076 and L038 independently, using the *FindMarkers* function from the Seurat package. For L076, we compared progressive cancer cells from relapse 1 and relapse 2 against those from the initial diagnostic samples. For L038, we compared cancer cells with *TP53* loss in timepoint 1 samples against cancer cells without *TP53* loss in the initial diagnostic samples. Full results can be found in **Supplementary Table 8**.

### Transcriptional module enrichment analysis

The trisomy 21 - leukaemic gene module was composed of all genes derived from the above analysis. The *GATA1* - leukaemic gene module and ML-DS leukaemic gene module from the above analyses were further refined to retain only genes expressed in >= 20% more cells in the group with higher expression. The enrichment score for each gene module (composed of both up- and down-regulated genes) in single-cell RNAseq data was calculated using the function *AddModuleScore_UCell* from the R package UCell^73^ (v1.3.1). For the bulk transcriptome dataset, the enrichment score was computed using the function *simpleScore* from the R package singscore^74^ (v1.10.0).

### Phylogenetic reconstruction of progressive ML-DS

#### Somatic variants calling

We investigated the genetic evolution of relapse and refractory ML-DS in child L076 and child L038 respectively. For both cases, the genetic phylogenetic relation was reconstructed based on somatic single-nucleotide variants (SNVs, also referred to as substitutions). SNVs in both the diagnostic and TP1 WGS samples were called using the CaVEMan algorithm^75^ (v1.18.2), and short insertions/deletions were called using Pindel algorithm^76^ (v3.10.0) in an unmatched analysis of each sample against an *in silico* normal human reference genome. In addition to the inbuilt QC filters, we further applied a series of stringent filters to remove low quality variants and germline variants, following the detailed workflow outlined in Coorens *et al.*, 2019^44^. Briefly, we only retained variants with a high median alignment score of supporting reads (ASMD >= 140), and required that fewer than half of the reads were clipped (CLPM=0), as well as not falling within 10 base pair of a deletion or insertion called by Pindel. We then recounted across both samples the variant allele frequency of all substitutions with a cut-off for base quality of 25 and read mapping quality of 30. Variants were also filtered out if they were called in a region of consistently low depth across both samples (excluding copy number segments). Next, to filter out germline substitutions, we fitted a binomial distribution to the combined read counts across all samples per SNV site, and applied a one-sided exact binomial test to calculate the probability that the SNV is consistent with being a germline variant. For the remaining variants, we calculated the site-specific error rates by interrogating the same sites in a panel of normal blood samples, using a beta-binomial model derived from the Shearwater variant call^77^. Variants which were indistinguishable from the background error rates were removed. All variants which passed all filters were visually inspected using the genome browser Jbrowse^78^ to exclude further sequencing or mapping artefacts. The final list of somatic substitutions can be found in **Supplementary Table 6** for L076 and **Supplementary Table 7** for L038.

#### Single-base substitution signature analysis

The single-base substitution profile of each WGS sample from L076 and L038 was further deconvolved into linear combinations of known COSMIC reference signatures (v3.4) using the R package mSigAct^79^ (v3.0.1). The set of reference signatures provided was: SBS1, SBS5, SBS13, SBS15, SBS18, SBS31, SBS35, SBS44, SBS84, and SBS86. Signature exposures per sample can be found in **Figure 4A and C**.

#### WGS-derived copy number aberrations

We identified copy number aberrations in each WGS sample from L076 and L038 using the COBALT, AMBER, and PURPLE toolkit developed by the Hartwig Medical Foundation, following their documentation available on github:https://github.com/hartwigmedical/hmftools. Briefly, the B-allele frequency (BAF) of heterozygous single-nucleotide polymorphism (SNP) sites was computed using AMBER (v3.9), and read depth ratios were calculated with COBALT (v1.14.1), both in “tumor-only” mode. B-allele frequency information and read-depth ratios were integrated to estimate the purity, ploidy and copy number profile of each tumour using PURPLE^80^ (v3.8.4).

#### Detection of copy number aberrations in scRNA-seq data

Using the R package alleleIntegrator^45^ (v0.9.1), we examined the copy number status of individual leukaemic blasts from patient L076 (relapse ML-DS), L038 (refractory ML-DS) and L067 (diploid MDS with *GATA1* mutation). Heterozygous SNPs were identified from WGS data from each individual, and phasing of heterozygous SNPs across copy number segments with allelic imbalance was performed. The number of scRNA-seq reads supporting the major/minor allele across each copy number segment in each cell was calculated. Finally, the posterior probability of each cell harbouring the altered or normal copy number state for each copy number segment was calculated.

## Reference

1. Labuhn, M. et al. Mechanisms of Progression of Myeloid Preleukemia to Transformed Myeloid Leukemia in Children with Down Syndrome. Cancer Cell 36, 123–138.e10 (2019).

2. Hasle, H., Clemmensen, I. H. & Mikkelsen, M. Risks of leukaemia and solid tumours in individuals with Down’s syndrome. The Lancet 355, 165–169 (2000).

3. Marlow, E. C. et al. Leukemia Risk in a Cohort of 3.9 Million Children with and without Down Syndrome. J. Pediatr. 234, 172–180.e3 (2021).

4. Roberts, I. et al. GATA1-mutant clones are frequent and often unsuspected in babies with Down syndrome: identification of a population at risk of leukemia. Blood 122, 3908–3917 (2013).

5. Goemans, B. F. et al. Sensitive GATA1 mutation screening reliably identifies neonates with Down syndrome at risk for myeloid leukemia. Leukemia 35, 2403–2406 (2021).

6. van den Akker, T. A., et al. Myeloid Proliferations Associated with Down Syndrome: Clinicopathologic Characteristics of Forty Cases from Five Large Academic Institutions. Pathobiology 91, 89–98 (2024).

7. Karandikar, N. J., Aquino, D. B., McKenna, R. W. & Kroft, S. H. Transient Myeloproliferative Disorder and Acute Myeloid Leukemia in Down Syndrome: An Immunophenotypic Analysis. Am. J. Clin. Pathol. 116, 204–210 (2001).

8. Bhatnagar, N., Nizery, L., Tunstall, O., Vyas, P. & Roberts, I. Transient Abnormal Myelopoiesis and AML in Down Syndrome: an Update. Curr. Hematol. Malig. Rep. 11, 333–341 (2016).

9. Tunstall, O. et al. Guidelines for the investigation and management of Transient Leukaemia of Down Syndrome. Br. J. Haematol. 182, 200–211 (2018).

10. Roy, A. et al. Perturbation of fetal liver hematopoietic stem and progenitor cell development by trisomy 21. Proc. Natl. Acad. Sci. 109, 17579–17584 (2012).

11. Popescu, D.-M. et al. Decoding human fetal liver haematopoiesis. Nature 574, 365–371 (2019).

12. Jardine, L. et al. Blood and immune development in human fetal bone marrow and Down syndrome. Nature 598, 327–331 (2021).

13. Marderstein, A. R. et al. Single-cell multi-omics map of human foetal blood in Down’s Syndrome. 2023.09.25.559431 Preprint at 10.1101/2023.09.25.559431 (2023).

14. Wechsler, J. et al. Acquired mutations in GATA1 in the megakaryoblastic leukemia of Down syndrome. Nat. Genet. 32, 148–152 (2002).

15. Greene, M. E. et al. Mutations in *GATA1* in both transient myeloproliferative disorder and acute megakaryoblastic leukemia of Down syndrome. Blood Cells. Mol. Dis. 31, 351–356 (2003).

16. Baruchel, A. et al. Down syndrome and leukemia: from basic mechanisms to clinical advances. Haematologica 108, 2570–2581 (2023).

17. Kuhl, C. et al. GATA1-Mediated Megakaryocyte Differentiation and Growth Control Can Be Uncoupled and Mapped to Different Domains in GATA1. Mol. Cell. Biol. 25, 8592–8606 (2005).

18. Welch, J. J. et al. Global regulation of erythroid gene expression by transcription factor GATA-1. Blood 104, 3136–3147 (2004).

19. Muntean, A. G. & Crispino, J. D. Differential requirements for the activation domain and FOG-interaction surface of GATA-1 in megakaryocyte gene expression and development. Blood 106, 1223–1231 (2005).

20. Bourquin, J.-P. et al. Identification of distinct molecular phenotypes in acute megakaryoblastic leukemia by gene expression profiling. Proc. Natl. Acad. Sci. 103, 3339–3344 (2006).

21. Sato, T. et al. Landscape of driver mutations and their clinical effects on Down syndrome–related myeloid neoplasms. Blood 143, 2627–2643 (2024).

22. Grimm, J., Heckl, D. & Klusmann, J.-H. Molecular Mechanisms of the Genetic Predisposition to Acute Megakaryoblastic Leukemia in Infants With Down Syndrome. Front. Oncol. 11, (2021).

23. Roy, A., Roberts, I. & Vyas, P. Biology and management of transient abnormal myelopoiesis (TAM) in children with Down syndrome. Semin. Fetal. Neonatal Med. 17, 196–201 (2012).

24. Köhnke, T. et al. Integrated multiomic approach for identification of novel immunotherapeutic targets in AML. Biomark. Res. 10, 43 (2022).

25. Miyauchi, M. et al. ADAM8 Is an Antigen of Tyrosine Kinase Inhibitor-Resistant Chronic Myeloid Leukemia Cells Identified by Patient-Derived Induced Pluripotent Stem Cells. Stem Cell Rep. 10, 1115–1130 (2018).

26. Evers, M. et al. Association of ADAM family members with proliferation signaling and disease progression in multiple myeloma. Blood Cancer J. 14, 1–11 (2024).

27. Debili, N. et al. Different expression of CD41 on human lymphoid and myeloid progenitors from adults and neonates. Blood 97, 2023–2030 (2001).

28. Nagata, Y. & Suzuki, R. FcεRI: A Master Regulator of Mast Cell Functions. Cells 11, 622 (2022).

29. Rönnberg, E. et al. Analysis of human lung mast cells by single cell RNA sequencing. Front. Immunol. 14, (2023).

30. Song, S.-H., Kim, A., Dale, R. & Dean, A. Ldb1 regulates carbonic anhydrase 1 during erythroid differentiation. Biochim. Biophys. Acta BBA - Gene Regul. Mech. 1819, 885–891 (2012).

31. Ramos-Mejía, V. et al. HOXA9 promotes hematopoietic commitment of human embryonic stem cells. Blood 124, 3065–3075 (2014).

32. Lee, J.-W. et al. DACH1 regulates cell cycle progression of myeloid cells through the control of cyclin D, Cdk 4/6 and p21Cip1. Biochem. Biophys. Res. Commun. 420, 91–95 (2012).

33. Wagenblast, E. et al. Mapping the cellular origin and early evolution of leukemia in Down syndrome. Science 373, eabf6202 (2021).

34. Hu, G. et al. PRMT2 accelerates tumorigenesis of hepatocellular carcinoma by activating Bcl2 via histone H3R8 methylation. Exp. Cell Res. 394, 112152 (2020).

35. Karagiota, A., Chachami, G. & Paraskeva, E. Lipid Metabolism in Cancer: The Role of Acylglycerolphosphate Acyltransferases (AGPATs). Cancers 14, 228 (2022).

36. Nagel, S. The Role of IRX Homeobox Genes in Hematopoietic Progenitors and Leukemia. Genes 14, 297 (2023).

37. Byrska-Bishop, M. et al. Pluripotent stem cells reveal erythroid-specific activities of the GATA1 N-terminus. J. Clin. Invest. 125, 993–1005 (2015).

38. Birger, Y. et al. Perturbation of fetal hematopoiesis in a mouse model of Down syndrome’s transient myeloproliferative disorder. Blood 122, 988–998 (2013).

39. Li, Z. et al. Developmental stage–selective effect of somatically mutated leukemogenic transcription factor GATA1. Nat. Genet. 37, 613–619 (2005).

40. Alejo-Valle, O. et al. The megakaryocytic transcription factor ARID3A suppresses leukemia pathogenesis. Blood 139, 651–665 (2022).

41. Gonzales, F. et al. Tetraspanin CD81 Supports Cancer Stem Cell Function and Represents a Therapeutic Vulnerability in Acute Myeloid Leukemia. Blood 142, 168 (2023).

42. Yu, H. et al. Bone marrow stromal cell antigen 2: Tumor biology, signaling pathway and therapeutic targeting (Review). Oncol. Rep. 51, 1–12 (2024).

43. Weisser, M. et al. PTK2 expression and immunochemotherapy outcome in chronic lymphocytic leukemia. Blood 124, 420–425 (2014).

44. Coorens, T. H. H. et al. Embryonal precursors of Wilms tumor. Science 366, 1247–1251 (2019).

45. Trinh, M. K. et al. Precise identification of cancer cells from allelic imbalances in single cell transcriptomes. *Commun*. Biol. 5, 1–8 (2022).

46. Qian, Z. et al. Enhanced expression of FHL2 leads to abnormal myelopoiesis in vivo. Leukemia 23, 1650–1657 (2009).

47. Li, Y. et al. CXCL8 is associated with the recurrence of patients with acute myeloid leukemia and cell proliferation in leukemia cell lines. Biochem. Biophys. Res. Commun. 499, 524–530 (2018).

48. He, Y. et al. Overexpression of *EPS8* is associated with poor prognosis in patients with acute lymphoblastic leukemia. Leuk. Res. 39, 575–581 (2015).

49. Ji, H. et al. CD82 supports survival of childhood acute myeloid leukemia cells via activation of Wnt/β-catenin signaling pathway. Pediatr. Res. 85, 1024–1031 (2019).

50. Floren, M. et al. Tetraspanin CD82 drives acute myeloid leukemia chemoresistance by modulating protein kinase C alpha and β1 integrin activation. Oncogene 39, 3910–3925 (2020).

51. Barwe, S. P., Sidhu, I., Kolb, E. A. & Gopalakrishnapillai, A. Modeling Transient Abnormal Myelopoiesis Using Induced Pluripotent Stem Cells and CRISPR/Cas9 Technology. Mol. Ther. Methods Clin. Dev. 19, 201–209 (2020).

52. Ling, T., Zhang, K., Yang, J., Gurbuxani, S. & Crispino, J. D. Gata1s mutant mice display persistent defects in the erythroid lineage. Blood Adv. 7, 3253–3264 (2023).

53. Arkoun, B. et al. Stepwise GATA1 and SMC3 mutations alter megakaryocyte differentiation in a Down syndrome leukemia model. J. Clin. Invest. 132, (2023).

54. Gialesaki, S. et al. RUNX1 isoform disequilibrium promotes the development of trisomy 21– associated myeloid leukemia. Blood 141, 1105–1118 (2023).

55. Lukes, J. et al. Chromosome 21 gain is dispensable for transient myeloproliferative disorder driven by a novel GATA1 mutation. Leukemia 34, 2503–2508 (2020).

56. Klusmann, J.-H. et al. Treatment and prognostic impact of transient leukemia in neonates with Down syndrome. Blood 111, 2991–2998 (2008).

57. Flasinski, M. et al. Low-dose cytarabine to prevent myeloid leukemia in children with Down syndrome: TMD Prevention 2007 study. Blood Adv. 2, 1532–1540 (2018).

58. Yamato, G. et al. Predictive factors for the development of leukemia in patients with transient abnormal myelopoiesis and Down syndrome. Leukemia 35, 1480–1484 (2021).

59. Alford, K. A. et al. Analysis of GATA1 mutations in Down syndrome transient myeloproliferative disorder and myeloid leukemia. Blood 118, 2222–2238 (2011).

60. Gerrelli, D., Lisgo, S., Copp, A. J. & Lindsay, S. Enabling research with human embryonic and fetal tissue resources. Development 142, 3073–3076 (2015).

61. Zheng, G. X. Y. et al. Massively parallel digital transcriptional profiling of single cells. Nat. Commun. 8, 14049 (2017).

62. Hao, Y. et al. Integrated analysis of multimodal single-cell data. Cell 184, 3573–3587.e29 (2021).

63. Wolock, S. L., Lopez, R. & Klein, A. M. Scrublet: Computational Identification of Cell Doublets in Single-Cell Transcriptomic Data. Cell Syst. 8, 281–291.e9 (2019).

64. Young, M. D. & Behjati, S. SoupX removes ambient RNA contamination from droplet-based single-cell RNA sequencing data. GigaScience 9, giaa151 (2020).

65. Chen, S., Zhou, Y., Chen, Y. & Gu, J. fastp: an ultra-fast all-in-one FASTQ preprocessor. Bioinforma. Oxf. Engl. 34, i884–i890 (2018).

66. Dobin, A. et al. STAR: ultrafast universal RNA-seq aligner. Bioinforma. Oxf. Engl. 29, 15–21 (2013).

67. Patro, R., Duggal, G., Love, M. I., Irizarry, R. A. & Kingsford, C. Salmon provides fast and bias-aware quantification of transcript expression. Nat. Methods 14, 417–419 (2017).

68. Li, H. & Durbin, R. Fast and accurate short read alignment with Burrows–Wheeler transform. Bioinformatics 25, 1754–1760 (2009).

69. Domínguez Conde, C., et al. Cross-tissue immune cell analysis reveals tissue-specific features in humans. Science 376, eabl5197 (2022).

70. Young, M. D. et al. Single-cell transcriptomes from human kidneys reveal the cellular identity of renal tumors. Science 361, 594–599 (2018).

71. Li, H. et al. The Sequence Alignment/Map format and SAMtools. Bioinformatics 25, 2078–2079 (2009).

72. Love, M. I., Huber, W. & Anders, S. Moderated estimation of fold change and dispersion for RNA-seq data with DESeq2. Genome Biol. 15, 550 (2014).

73. Andreatta, M. & Carmona, S. J. UCell: Robust and scalable single-cell gene signature scoring. Comput. Struct. Biotechnol. J. 19, 3796–3798 (2021).

74. Foroutan, M. et al. Single sample scoring of molecular phenotypes. BMC Bioinformatics 19, 404 (2018).

75. Jones, D. et al. cgpCaVEManWrapper: Simple Execution of CaVEMan in Order to Detect Somatic Single Nucleotide Variants in NGS Data. Curr. Protoc. Bioinforma. 56, 15.10.1–15.10.18 (2016).

76. Ye, K. et al. Split-Read Indel and Structural Variant Calling Using PINDEL. in Copy Number Variants: Methods and Protocols (ed. Bickhart, D. M.) 95–105 (Springer New York, New York, NY, 2018). doi:10.1007/978-1-4939-8666-8_7.

77. Gerstung, M., Papaemmanuil, E. & Campbell, P. J. Subclonal variant calling with multiple samples and prior knowledge. Bioinformatics 30, 1198–1204 (2014).

78. Diesh, C. et al. JBrowse 2: a modular genome browser with views of synteny and structural variation. Genome Biol. 24, 74 (2023).

79. Ng, A. W. T. et al. Aristolochic acids and their derivatives are widely implicated in liver cancers in Taiwan and throughout Asia. Sci. Transl. Med. 9, eaan6446 (2017).

80. Priestley, P. et al. Pan-cancer whole-genome analyses of metastatic solid tumours. Nature 575, 210–216 (2019).

