## Supplemental Figures for "Single cell transcriptional evolution of myeloid leukaemia of Down syndrome"

**Affiliations**:

*Co-corresponding and co-directing authors

Contact information:

### Supplementary Tables

**Supplementary Table 1**: Paediatric leukaemia and fetal liver datasets overview.

**Supplementary Table 2**: Differential gene expression analysis comparing cells from fetuses with abnormal karyotypes against diploid cells, for each cell type and each karyotype.

**Supplementary Table 3**: List of differentially expressed genes representing the contribution of trisomy 21 towards leukaemic transcriptome.

**Supplementary Table 4**: List of differentially expressed genes representing the transcriptomic consequences of *GATA1* mutations towards leukaemic transcriptome.

**Supplementary Table 5**: List of differentially expressed genes representing the transcriptomic differences between conventional TAM blasts and ML-DS diagnostic bone marrow blasts.

**Supplementary Table 6**: Catalogue of somatic mutations in L076 with relapse ML-DS.

**Supplementary Table 7**: Catalogue of somatic mutations in L038 with refractory ML-DS.

**Supplementary Table 8**: List of markers when comparing progressive ML-DS blasts against diagnostic ML-DS blasts from the same children, or all other responsive ML-DS blasts.

### Supplementary Figure Legends

### Supplementary Figure 1

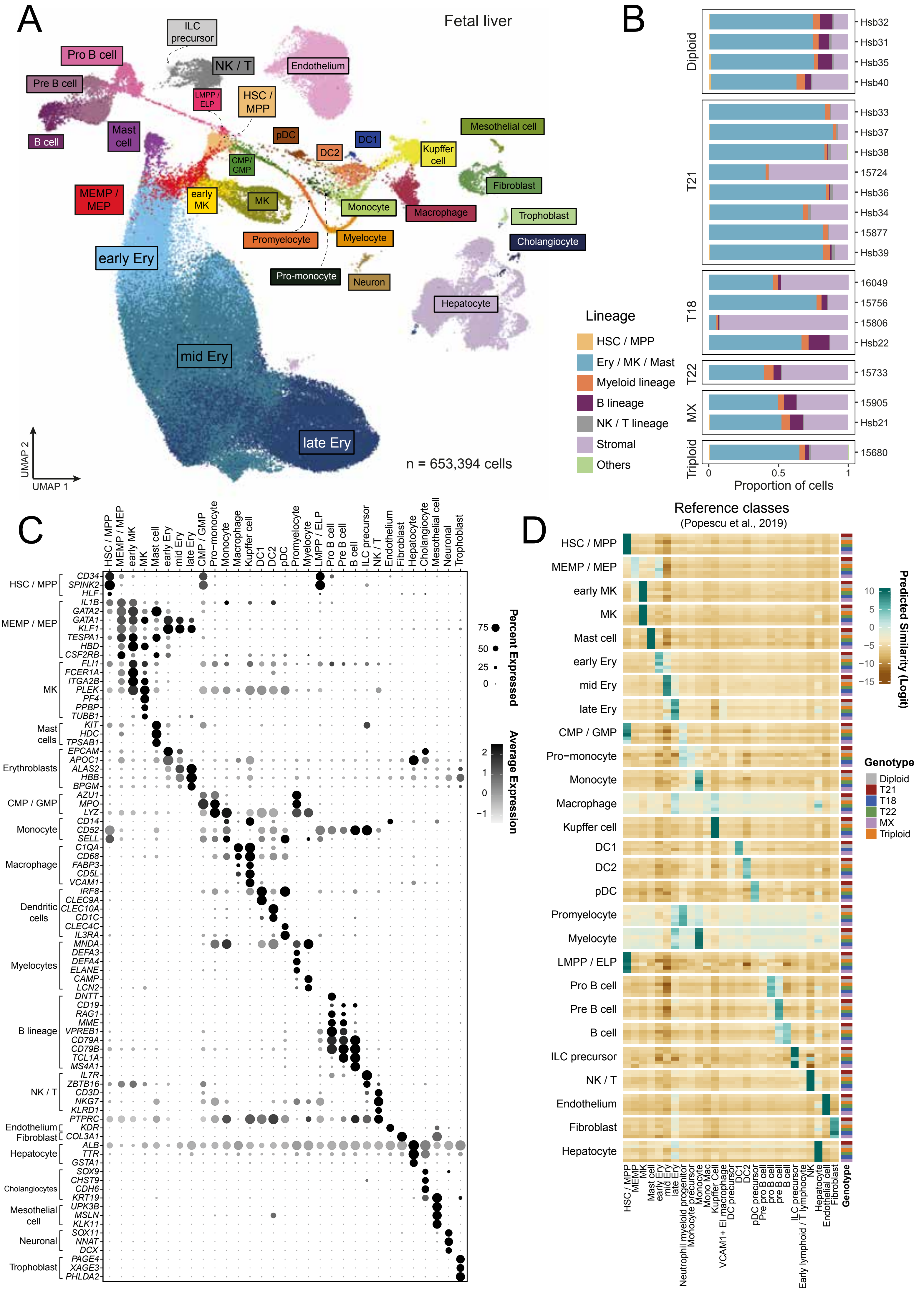

**Supplementary Figure 1: Fetal liver scRNA-seq dataset.**

(A) Uniform Manifold Approximation and Projection (UMAP) visualisation of the fetal liver scRNA-seq dataset (detailed in **Table 1, Supplementary Table 1**), where cells (dots) are coloured by cell types.

(B) Bar plot showing the proportions of different lineages captured from each fetal liver sample.

(C) Dot plot showing the z-scaled mean expression levels (colour) of key cell-type defining marker genes. Dot size represents the proportion of cells within each category with positive expression.

(D) Heatmap showing the average predicted similarity score (calculated using a CellTypist logistic regression model) for each query cell type from our fetal liver scRNA-seq dataset (row panels) compared to reference cell types from the published cellular fetal liver atlas by Popescu, DM., *et al.*, 2019[11] (columns). Within each query cell type, cells are further grouped based on their respective karyotypes (individual rows). Darker green indicates stronger similarity, darker brown indicates stronger dissimilarity.

***Abbreviation***

Cell types: CMP / GMP - common myeloid progenitor / granulocyte-monocyte progenitor; DC - dendritic cell; Ery - erythroblast; HSC / MPP - haematopoietic stem cell / multipotent progenitor; ILC precursor - innate lymphoid cell precursor; LMPP / ELP - lymphoid-primed multipotent progenitor / early lymphoid progenitor; MEMP/MEP - megakaryocyte-erythroid-mast cell progenitor / megakaryocyte-erythroid progenitor; MK - megakaryocyte; Mono Mac - monocyte / macrophage; NK / T - natural killer cell / T cell; pDC - plasmacytoid dendritic cell; VCAM1+ EI macrophage - VCAM1+ erythroblastic island macrophages.

Genotypes: T - trisomy; MX - monosomy X.

**Supplementary Figure 2**

# A

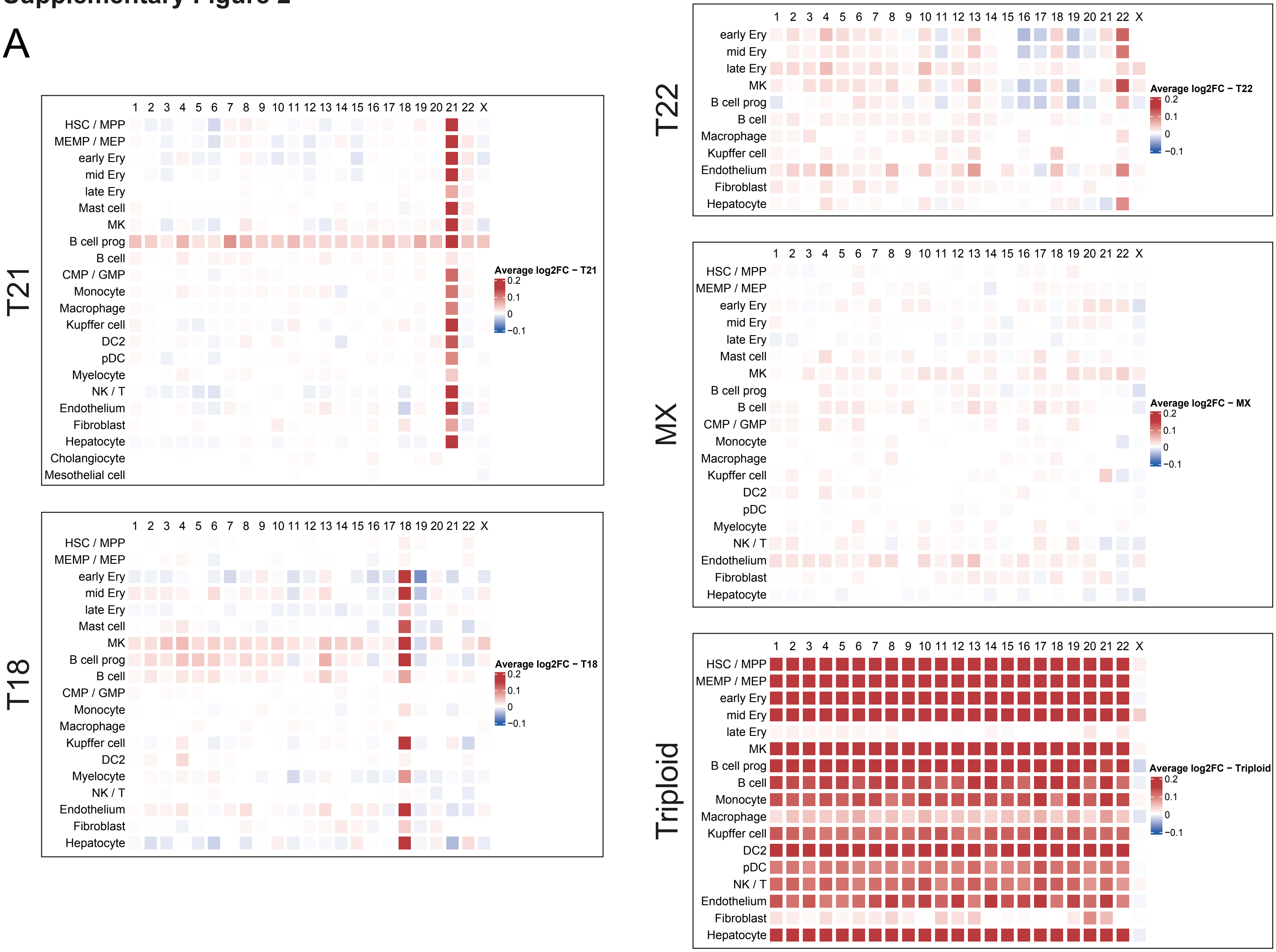

**Supplementary Figure 2: Transcriptional consequences of abnormal karyotypes across fetal hepatic cell types**

(A) Per-chromosome median log2 fold change of gene expression levels when comparing each abnormal karyotype against the corresponding diploid counterpart for each cell type. Darker red indicates higher expression level in cells with abnormal karyotype, darker blue indicates decreased expression in cells with abnormal karyotype. “B cell prog” includes progenitors of B-cell lineage: LMPP / ELP, pro B cell, and pre B cell; “NK / T” includes ILC precursors and NK / T cells.

***Abbreviation***

Cell types: CMP / GMP - common myeloid progenitor / granulocyte-monocyte progenitor;
DC - dendritic cell; Ery - erythroblast; HSC / MPP - haematopoietic stem cell / multipotent progenitor;
LMPP / ELP - lymphoid-primed multipotent progenitor / early lymphoid progenitor;
MEMP/MEP - megakaryocyte-erythroid-mast cell progenitor / megakaryocyte-erythroid progenitor;
MK - megakaryocyte; NK / T - natural killer cell / T cell; pDC - plasmacytoid dendritic cell.
Genotypes: T - trisomy; MX - monosomy X.

### Supplementary Figure 3

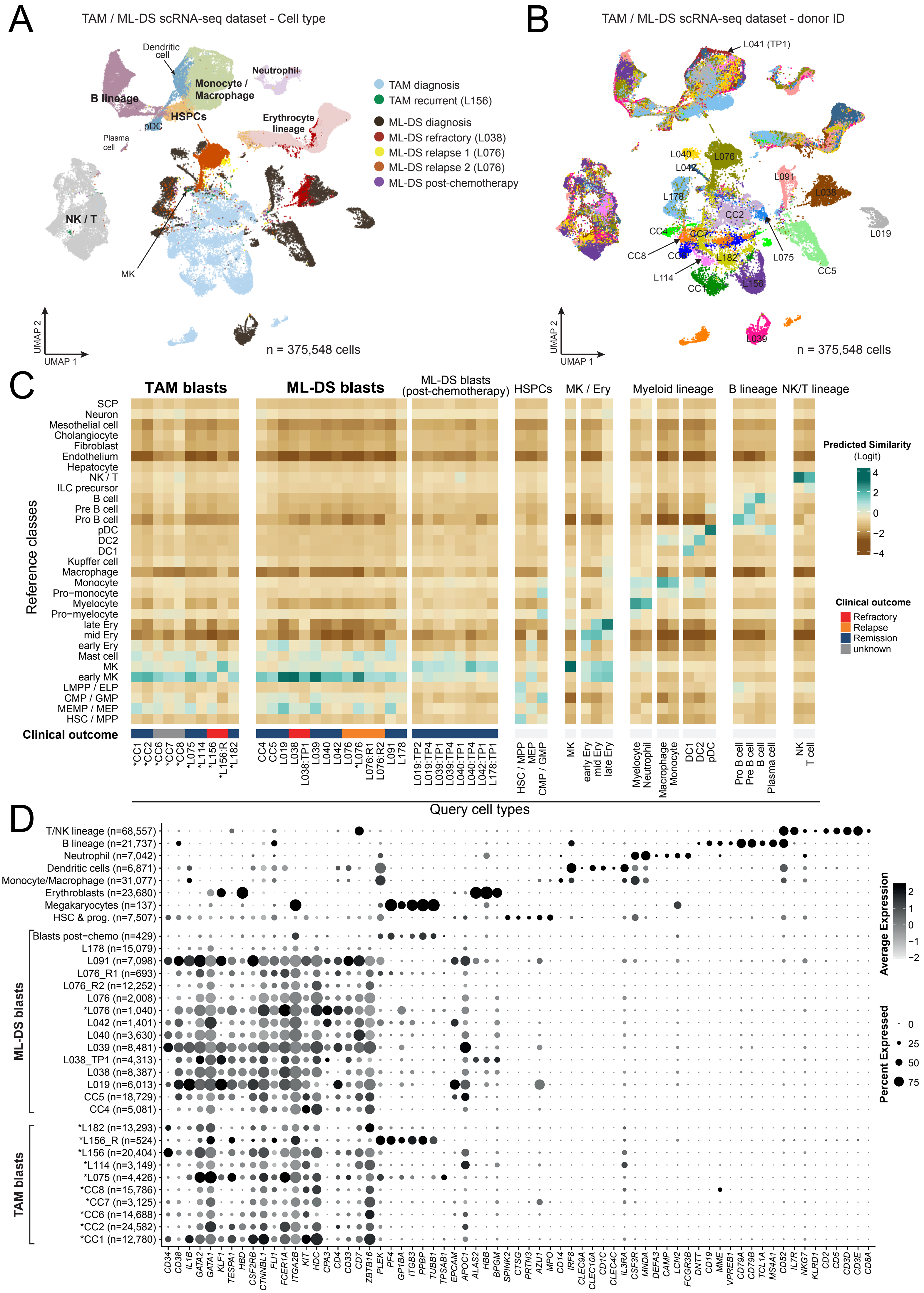

**Supplementary Figure 3: TAM / ML-DS scRNA-seq dataset.**

(A) UMAP visualisation of the TAM / ML-DS scRNA-seq dataset (detailed in **Table 1, Supplementary Table 1**), where cells (dots) are coloured by their corresponding cell types.

(B) UMAP visualisation of the TAM / ML-DS scRNA-seq dataset, where cells (dots) are coloured by the corresponding donor ID. Leukaemic blasts generally form donor-specific clusters, whereas normal cells across donors are clustered by cell type identities.

(C) Heatmap showing the average predicted similarity score (calculated by a logistic regression model) for each query cell type from our TAM / ML-DS scRNA-seq dataset (column panels) compared to reference cell types from our fetal liver scRNA-seq dataset (rows). Darker green indicates stronger similarity, darker brown indicates stronger dissimilarity. Leukaemic cells are grouped by patient / timepoint / tissue. Samples are bone marrow aspirates collected at initial diagnosis (treatment naive), unless indicated otherwise in the group name. Asterisks indicate samples from peripheral blood, otherwise are bone marrow aspirates. Leukaemic cells broadly resemble cell types along the erythrocyte-megakaryocyte-mast cell lineage, with the strongest similarity against the early megakaryocyte signal. Cell types from the normal compartment generally strongly match the corresponding reference classes. Asterisks (*) indicate peripheral blood samples; all others are bone marrow aspirates.

(D) Dot plot showing the z-scaled mean expression levels (colour) of key cell type – defining marker genes. Dot size represents the proportion of cells within each category with positive expression. Normal cells are grouped into their corresponding lineages as shown in (C). Leukaemic blasts are grouped by patient / timepoint / tissue, as detailed in (C). The number of cells in each category is indicated inside brackets. Asterisks indicate peripheral blood samples; all others are bone marrow aspirates.

***Abbreviation***

<ins>Cell types</ins>: HSPCs - haematopoietic stem and progenitor cells;
CMP / GMP - common myeloid progenitor / granulocyte-monocyte progenitor; DC - dendritic cell;
Ery - erythroblast; HSC / MPP - haematopoietic stem cell / multipotent progenitor;
ILC precursor - innate lymphoid cell precursor;
LMPP / ELP - lymphoid-primed multipotent progenitor / early lymphoid progenitor;
MEMP / MEP - megakaryocyte-erythroid-mast cell progenitor / megakaryocyte-erythroid progenitor;
MK - megakaryocyte; NK / T - natural killer cell / T cell; pDC - plasmacytoid dendritic cell;
SCP - schwann cell precursor.

<ins>Sample timepoint</ins>: R - recurrent; R1 - relapse 1 diagnosis; R2D - relapse 2 diagnosis;
TP1 - timepoint 1, TP2 - timepoint 2, TP4 - timepoint 4.

### Supplementary Figure 4

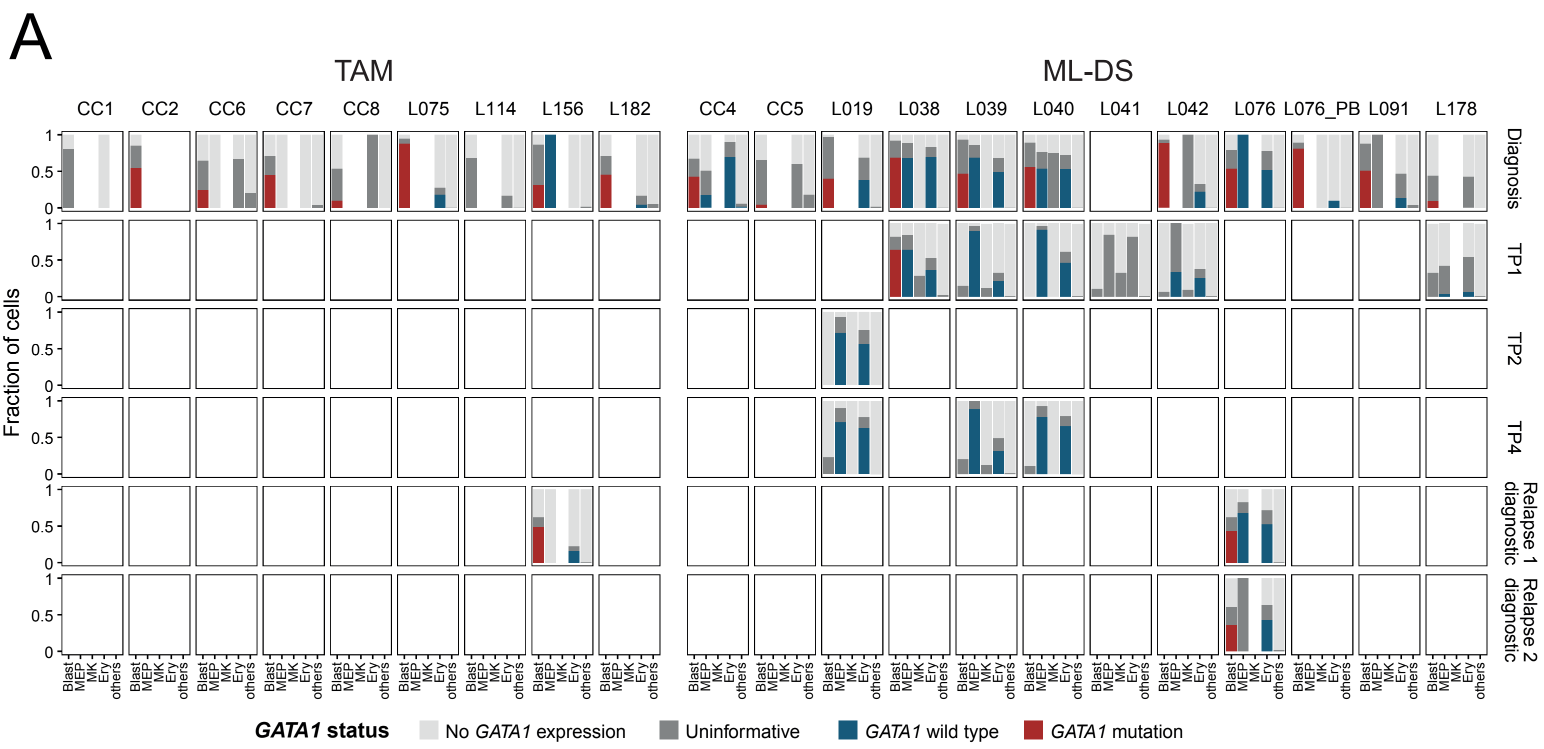

**Supplementary Figure 4: Single cell genotyping for *GATA1* mutation.**

(A) Bar plots showing the proportion of cells from each patient and timepoint categorised by *GATA1* genotypes. TAM/ML-DS blasts and cells along the megakaryocyte-erythroid-mast lineage are shown as individual columns; all other cells are grouped as "others" category. *GATA1* mutations are only detected in the blast populations. Asterisks (*) indicate peripheral blood samples; all others are bone marrow aspirates.

***Abbreviation***

<u>Sample timepoint</u>: TP1 - timepoint 1, TP2 - timepoint 2, TP4 - timepoint 4.

#### Supplementary Figure 5

A

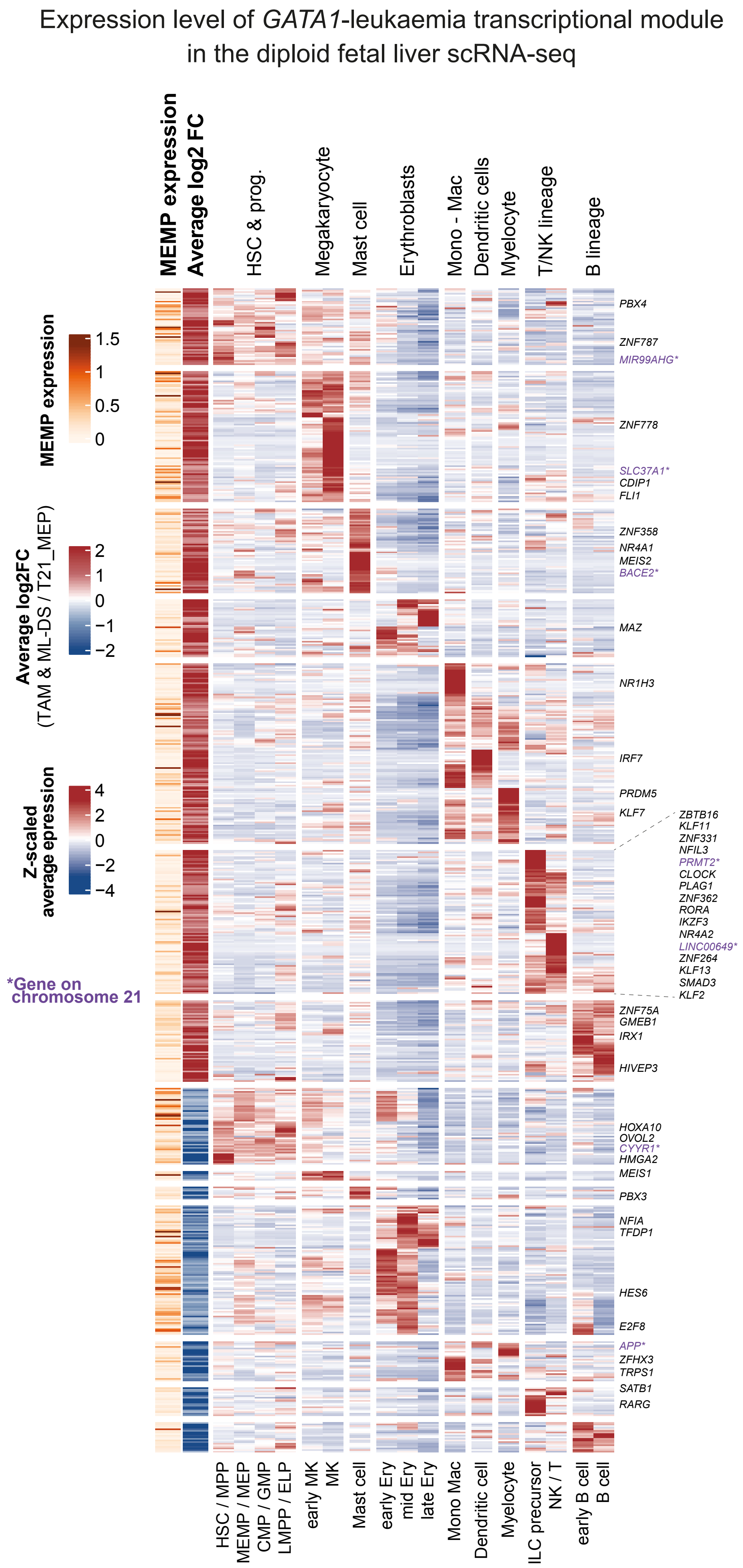


B

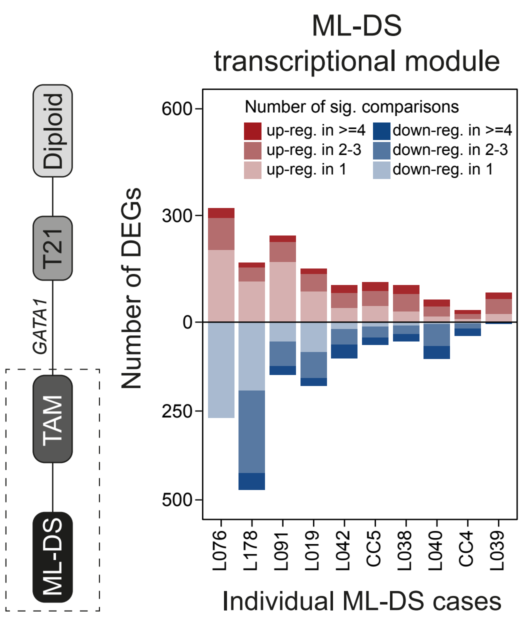

**Supplementary Figure 5: Transcriptional changes underpinning TAM and ML-DS.**

(A) Heatmap showing the z-scaled mean expression levels of the *GATA1* - leukaemia transcriptional module across haematopoietic lineages (columns) in our diploid fetal liver scRNA-seq. Darker red indicates higher expression level. Only genes expressed in the diploid fetal liver (see Methods) are included. “early B cell” includes pro B cell and pre B cell; “dendritic cells” include dendritic cell 1, dendritic cell 2, and plasmacytoid dendritic cell. Genes encoding for transcriptional factors are listed, genes located on chromosome 21 are listed and highlighted in purple.

(B) Bar plot showing the number of differentially expressed genes detected when comparing diagnostic (treatment naive) blasts from each ML-DS case (x-axis) against all conventional TAM blasts. Genes are further grouped based on the number of comparisons that they were identified as significantly expressed in. Darker red indicates up-regulated genes and darker blue indicates down-regulated genes which are detected in multiple ML-DS cases when compared to conventional TAM.

***Abbreviation***

Cell types: CMP / GMP - common myeloid progenitor / granulocyte-monocyte progenitor;
Ery - erythroblast; HSC / MPP - haematopoietic stem cell / multipotent progenitor;
ILC precursor - innate lymphoid cell precursor;
LMPP / ELP - lymphoid-primed multipotent progenitor / early lymphoid progenitor;
MEMP / MEP - megakaryocyte-erythroid-mast cell progenitor / megakaryocyte-erythroid progenitor;
MK - megakaryocyte; Mono/Mac - monocyte / macrophage; NK / T - natural killer cell / T cell.

### Supplementary Figure 6

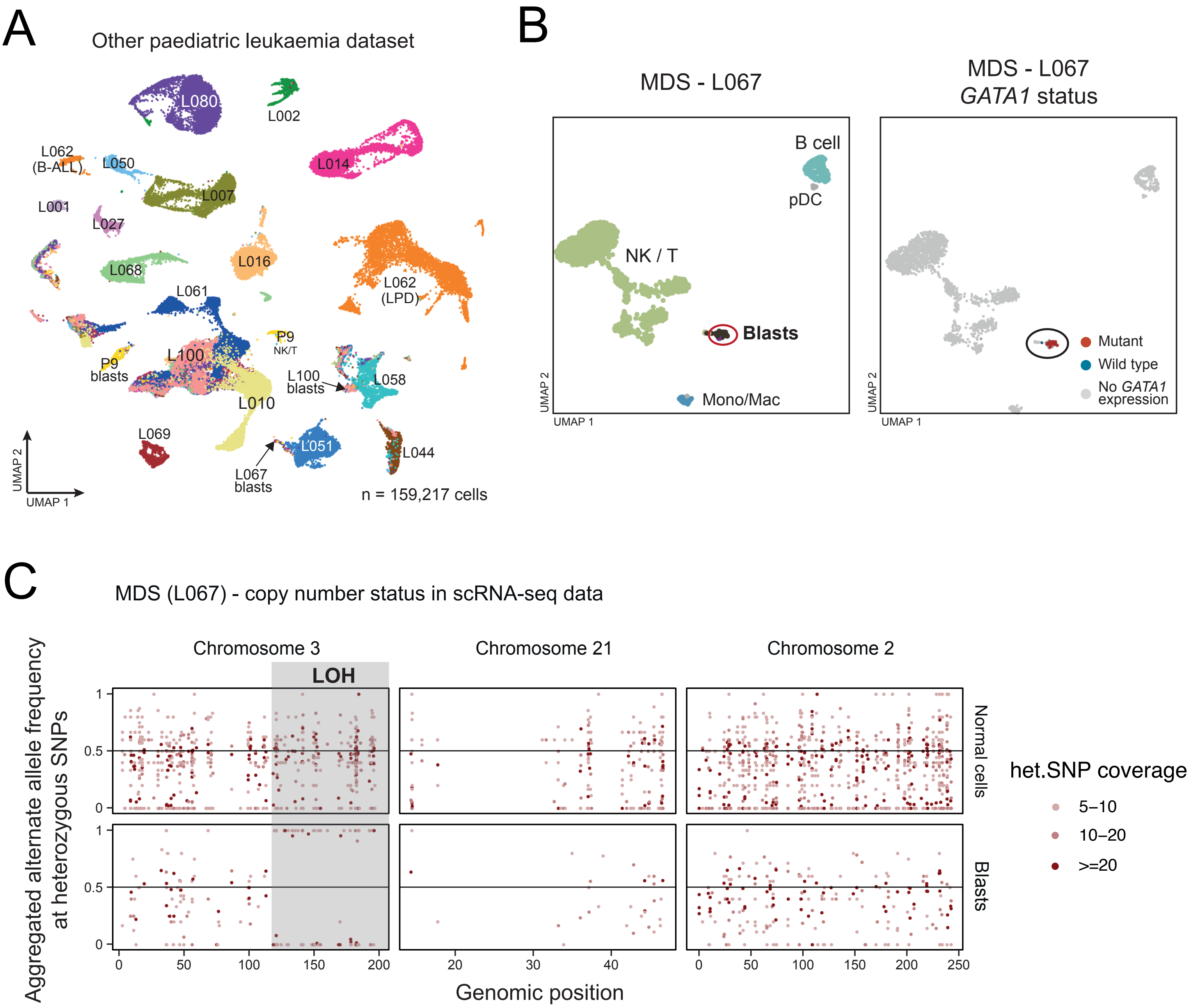

**Supplementary Figure 6: Other leukaemias scRNA-seq dataset.**

(A) UMAP visualisation of the additional scRNA-seq dataset of other leukaemias (detailed in **Table 1, Supplementary Table 1**), with cells (dots) coloured by donor ID. B-ALL - B cell acute lymphoblastic leukaemia; LPD - lymphoproliferative disorder.

(B) UMAP visualisation of the diploid myeloid neoplasm with *GATA1* mutation (MDS - myelodysplastic syndrome, donor L067). Cells (dots) are coloured by cell type (left) and the *GATA1* genotyping result status (right). pDC - plasmacytoid dendritic cell, NK / T - natural killer cell / T cell, Mono/Mac - monocyte / macrophage.

(C) Alternate-allele frequency (y-axis) of heterozygous single-nucleotide polymorphisms (het.SNPs) on chromosomes 3 (left), 21 (middle), and 2 (right) in scRNA-seq data of the diploid myeloid neoplasm with *GATA1* mutation (donor L067). The allele frequency is aggregated across blasts (bottom track) and normal cells (top track). Each dot represents a het.SNP on the chromosome, coloured by the total coverage across all cells in the group. Grey-shaded region highlights the copy number loss on chromosome 3, where loss-of-heterozygosity is observed in blasts but not in normal cells. There is no evidence of a shift in alternate allele frequency across het.SNPs on chromosome 21, thus no indication of copy number alteration on chromosome 21.

### Supplementary Figure 7

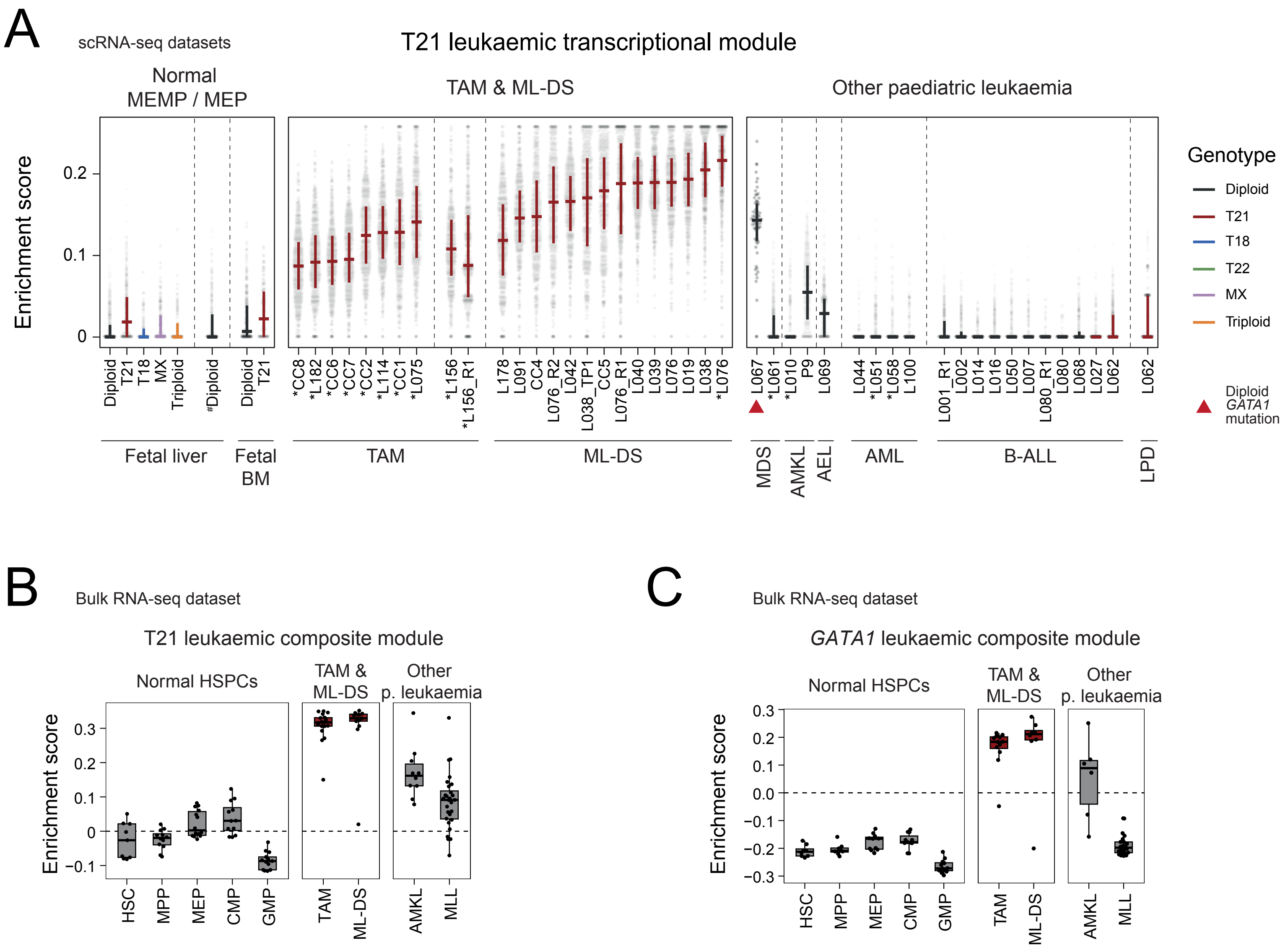

**Supplementary Figure 7: Specificity of the trisomy 21- and *GATA1* - leukaemia transcriptional modules.**

(A) Enrichment score of the trisomy 21 - leukaemia transcriptional module (y-axis) across individual cells (grey dots) from various scRNA-seq datasets (x-axis) consisting of: normal MEMP/MEP with different karyotypes from the fetal liver (including diploid cells from Popescu, D-M. *et al.*, 2019[11], denoted by #) and fetal bone marrow (BM) from Jardine, L. *et al.*, 2021[12]; TAM/ML-DS blasts; and leukaemic blasts from other leukaemias. Leukaemic cells are grouped by patient, timepoint (initial diagnosis unless indicated otherwise in the group name) and tissue. Amongst leukaemia samples, asterisks (*) indicate peripheral blood samples; all others are bone marrow aspirates. Cross bars indicate the interquartile range of enrichment score distribution.

(B) Enrichment score (y-axis) of the trisomy 21 - leukaemia transcriptional module in the independent bulk RNA-seq dataset composed of FACS-sorted normal haematopoietic stem and progenitor cells (HSPCs), TAM/ML-DS diagnostic samples, and leukaemic blasts from other paediatric leukaemias.

(C) Enrichment score (y-axis) of the *GATA1* - leukaemia transcriptional module (top genes) in the independent bulk RNA-seq dataset (as detailed in (B)).

***Abbreviation***

Leukaemia: MDS - myelodysplastic syndrome; AMKL - acute megakaryoblastic leukaemia; AML - acute myeloid leukemia; AEL - acute erythroid leukemia; B-ALL - B cell acute lymphoblastic leukaemia; LPD - lymphoproliferative disorder; MLL - mixed lineage leukaemia.

Cell types: HSPCs - haematopoietic stem and progenitor cells; HSC - haematopoietic stem cell; MPP - multipotent progenitor; MEP - megakaryocyte-erythroid progenitor; CMP - common myeloid progenitor; GMP - granulocyte-monocyte progenitor; NK / T - natural killer cell / T cell.

Genotypes: T - trisomy; MX - monosomy X.

Sample timepoint: R - recurrent; R1 - relapse 1 diagnosis; R2D - relapse 2 diagnosis; TP1 - timepoint 1.

### Supplementary Figure 8

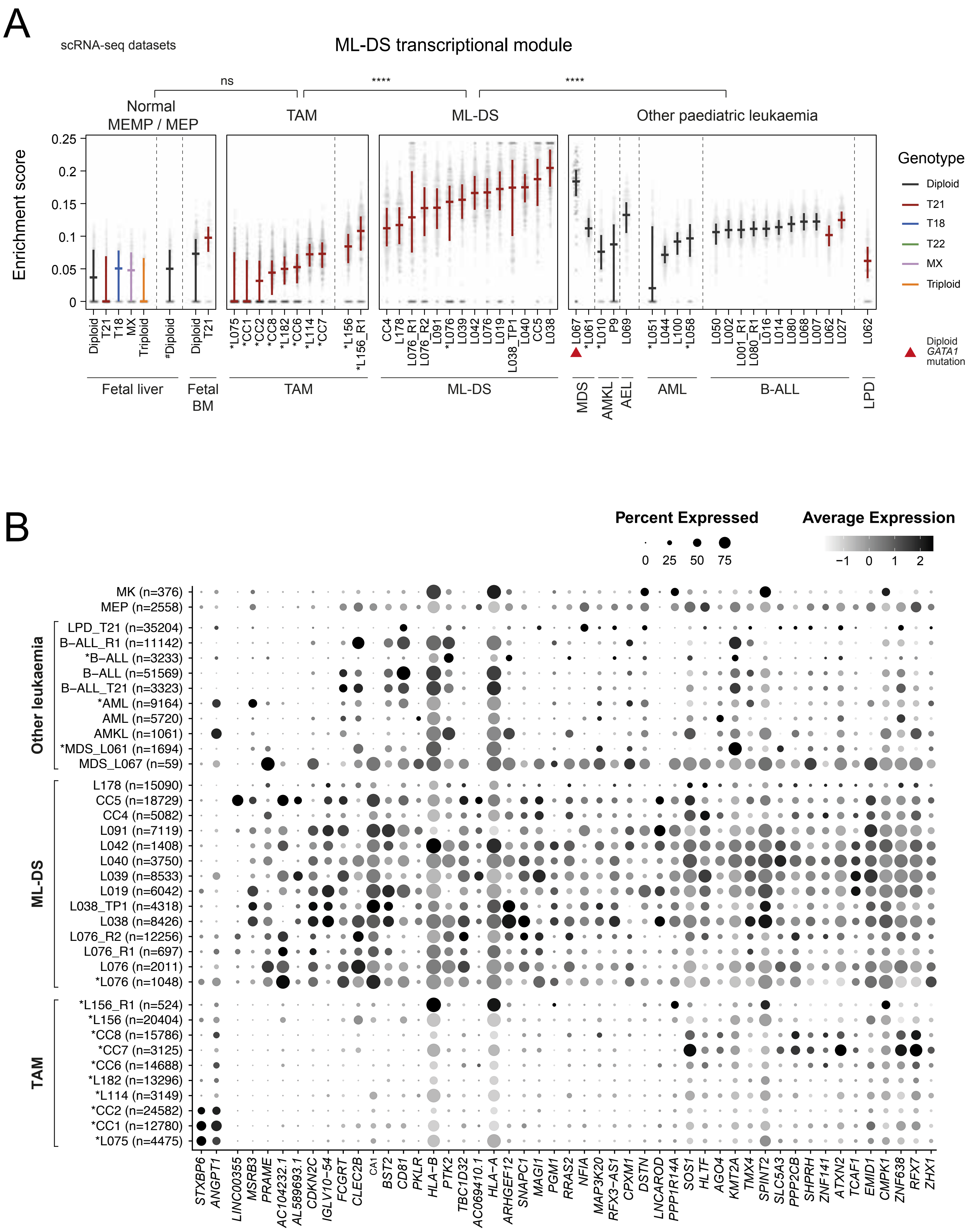

**Supplementary Figure 8: Specificity of the ML-DS leukaemia transcriptional module.**

(A) Enrichment score (y-axis) of the ML-DS transcriptional signature across individual cells (grey dots) from different scRNA-seq datasets (x-axis) consisting of: normal MEMP/MEP with different karyotypes from the fetal liver (including diploid cells from Popescu, D-M. *et al.*, 2019[11], denoted by #) and fetal bone marrow (BM) from Jardine, L. *et al.*, 2021[12]; TAM/ML-DS blasts; and leukaemic blasts from other leukaemias. Leukaemic cells are grouped by patient, timepoint (initial diagnosis unless indicated otherwise in the group name) and tissue. Amongst leukaemia samples, asterisks (*) indicate peripheral blood samples; all others are bone marrow aspirates. Cross bars indicate the interquartile range of enrichment score distribution.

(B) Dot plot showing z-scaled mean expression levels (colour) of top up- and down-regulated genes in ML-DS diagnostic blasts compared to conventional TAM blasts (i.e. the ML-DS leukaemic gene module). Dot size represents the proportion of cells with positive expression. Leukaemic cells are grouped by subtype, patient, timepoint (initial diagnosis unless indicated otherwise in the group name) and tissue. Asterisks indicate peripheral blood samples; all others are bone marrow aspirates. Normal megakaryocyte-erythrocyte progenitors (MEP) and megakaryocytes (MK) obtained from both leukaemia datasets are also shown. The number of cells in each group is indicated inside brackets (the *n* values).

***Abbreviation***

Cell types: MEP - megakaryocyte-erythroid progenitor; MK - megakaryocyte.
Leukaemia: MDS - myelodysplastic syndrome; AMKL - acute megakaryoblastic leukaemia;
AML - acute myeloid leukemia; AEL - acute erythroid leukemia;
B-ALL - B cell acute lymphoblastic leukaemia; LPD - lymphoproliferative disorder.
Sample timepoint: R - recurrent; R1 - relapse 1 diagnosis; R2D - relapse 2 diagnosis;
TP1 - timepoint 1, TP2 - timepoint 2, TP4 - timepoint 4.

### Supplementary Figure 9

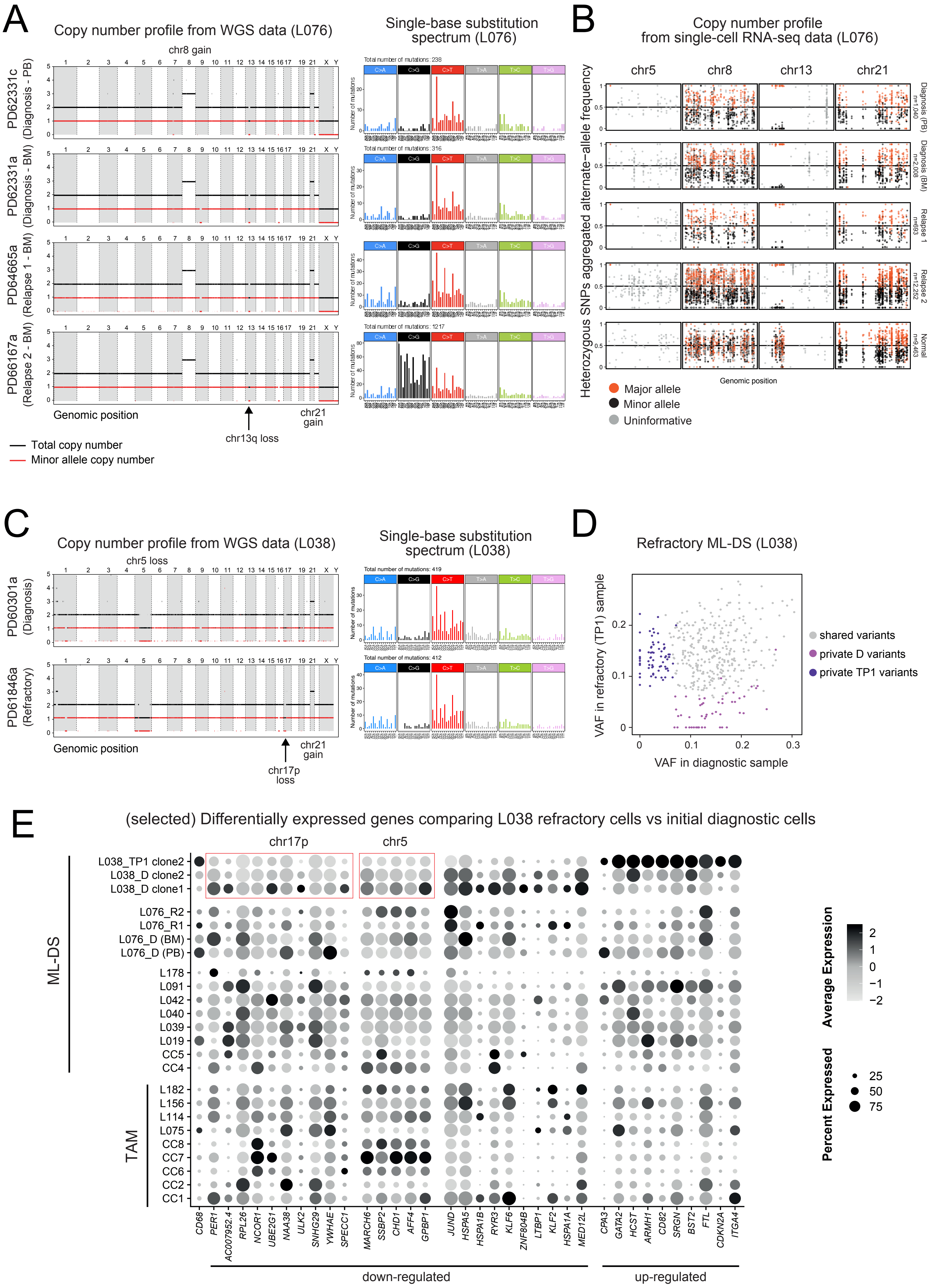

**Supplementary Figure 9: The genetic and transcriptional evolution of progressive ML-DS.**

(A) Copy number profiles derived from whole genome sequencing (WGS) data four samples from child L076 with two relapses after initial treatment for ML-DS; as called by HMF-PURPLE. Note that PURPLE is unable to determine subclonal copy number aberrations. For each chromosome, the black line indicates the total copy number, and the red line indicates the minor allele copy number. All samples showed chromosome 21 gain, chromosome 8 gain, and chromosome 13q loss.

(B) Copy number genotyping in scRNA-seq data from the same samples as in (A) from child L076. Cells are grouped by identity (normal or cancer) and sample timepoint (rows). The aggregated B-allele frequency (BAF) (y-axis) of heterozygous single-nucleotide polymorphisms (SNPs) across chromosomes 5, 8, 13 and 21 (columns) is shown for each cell category. Each dot represents a heterozygous SNP. Orange dots indicate alternate alleles located on the major chromosome, whereas black dots represent alternate alleles located on the minor chromosome. Grey dots are heterozygous SNPs with insufficient evidence of BAF deviation from the expected value of 0.5. Unlike normal cells, cancer cells from all timepoints exhibit deviation in BAF of heterozygous SNPs on chromosome 8 and 21, as well as chromosome 13q (loss of heterozygosity). This confirms the identity of the cancer cells, and suggests that these copy number alterations are early events, thus pervading all cancer cells.

(C) Copy number profiles from WGS data of two samples from child L038 with refractory ML-DS; analysed using HMF-PURPLE. Note that PURPLE is unable to determine whether the copy number aberrations are subclonal. For each chromosome, the black line indicates the total copy number, and the red line indicates the minor allele copy number. All samples showed chromosome 21 gain and 17p loss. The diagnostic sample showed evidence of chromosome 5q loss, whereas the timepoint 1 sample showed loss of the entire chromosome 5.

(D) Scatter plot showing the observed variant allele frequency (VAF) distribution from WGS data of two samples from child L038: diagnostic sample on x-axis and TP1 sample on y-axis. D - Diagnostic; TP1 - timepoint 1 (post 1 course of chemotherapy).

(E) Dot plot showing the average expression of top differentially expressed genes in clone 2 from TP1 sample (with chr17p loss, including loss of *TP53*) compared to clone 1 from diagnostic sample (without chr17p loss). Dot size represents the percentage of cells expressing each gene; colour shows the z-scaled normalised expression levels, where darker grey indicates higher expression.

***Abbreviation***

Chr - chromosome

Sample timepoint: D - diagnosis; R1 - relapse diagnosis; R2 - relapse 2 diagnosis; TP1 - timepoint 1.

Tissue: PB - peripheral blood; BM - bone marrow.
